# Multicellular Aligned Bands Disrupt Global Collective Cell Behavior

**DOI:** 10.1101/2022.05.30.494066

**Authors:** Mahvash Jebeli, Samantha K. Lopez, Zachary E. Goldblatt, Dannel McCollum, Sebastian Mana-Capelli, Qi Wen, Kristen Billiar

**Author notes:** Correspondence to: Kristen L. Billiar, Biomedical Engineering Department, Worcester Polytechnic Institute, 100 Institute Road, Worcester, MA 01609.

## Abstract

Mechanical stress patterns emerging from collective cell behavior have been shown to play critical roles in morphogenesis, tissue repair, and cancer metastasis. In our previous work utilizing microcontact printing to geometrically constrain valvular interstitial cell monolayers into specific shapes, we demonstrated that the general patterns of observed cell alignment, size, and apoptosis correlate with predicted mechanical stress fields if nonuniform cell properties are used in the computational models. However, these radially symmetric models did not predict the substantial heterogeneity in cell behavior observed in individual circular aggregates. The goal of this study is to determine how the heterogeneities in cell behavior emerge over time and diverge from the predicted collective cell behavior. Cell-cell interactions in circular multicellular aggregates of valvular interstitial cells were studied with time-lapse imaging ranging from hours to days, and migration, proliferation, and traction stresses were measured. Our results indicate that individual elongated cells create strong local alignment within pre-confluent cell populations on the microcontact printed protein islands. These cells influence the alignment of additional cells to create dense, locally aligned bands of cells which disrupt the global behavior. Cells are highly elongated at the endpoints of the bands yet have decreased spread area in the middle and reduced mobility. Although traction stresses at the endpoints of bands are enhanced, even to the point of detaching aggregates from the culture surface, the cells in dense bands exhibit reduced proliferation, less nuclear YAP, and increased apoptotic rates indicating a low stress environment. These findings suggest that strong local cell-cell interactions between primary fibroblastic cells can disrupt the global collective cellular behavior leading to substantial heterogeneity of cell behaviors in constrained monolayers. This local emergent behavior within aggregated fibroblasts may play an important role in development and disease of connective tissues.

## 1. Introduction

Morphogenesis, tissue repair, and cancer metastasis are driven by collective cell behaviors which differ substantially from independent cell behavior. Studies that use voids, scratch assays (Trepat et al. 2011, Camley et al. 2017, Ladoux et al. 2017, Nanavati et al. 2020, Vishwakarma et al. 2020), and micropatterned protein “islands” (Streichan et al. 2014, He et al. 2015) in specific geometries (Nelson et al. 2005, Li et al. 2009, Trepat et al. 2009, Moussus et al. 2014, Ng et al. 2014) demonstrate that emergent mechanical stress patterns contribute in large part to the collective cell behavior. In particular, maximum traction stresses at the surface correlate positively with proliferation at pattern edges (Streichan et al. 2014), while the anisotropy of computed cell layer stresses correlate with cell alignment and elongation (Streichan et al. 2014). The majority of studies on collective cell behavior utilize epithelial cells and cell lines which exhibit relatively uniform cell area when in a monolayer as well as strong contact inhibition. Emergent patterns in the behavior of these cell types are explained relatively well with computational mechanical models that assume uniform cell mechanical properties such as modulus and contractility (Nelson et al. 2005, Li et al. 2009, Moussus et al. 2014, Paek et al. 2021). However, primary mesenchymal cells, which are more contractile and have strong cell-cell interactions, exhibit less uniform behavior when cultured as monolayers (Ladoux et al. 2017, Xie et al. 2021).

The complex collective cell behavior of primary valvular interstitial cells (VICs) is is implicated in calcific aortic valve disease (Yip Cindy Ying et al. 2009, Bogdanova et al. 2019). VICs cultured as monolayers detach under high tension and form multicellular aggregates which become hyperconfluent. The cells in these high-density regions undergo apoptosis and calcification (Jian et al. 2003, Yip Cindy Ying et al. 2009, Bogdanova et al. 2019). Previously, we used uniform circular protein islands to form consistent multicellular aggregates to study the effects of collective cell behavior on calcification of these cells (Cirka et al. 2017). We found that the general patterns of cell alignment, size, and apoptosis correlate with predicted mechanical stress fields if nonuniform cell properties are used in computational models (Goldblatt et al. 2020). However, these radially symmetric models did not predict the substantial heterogeneity in cell behavior observed in individual circular aggregates. In particular, we observed asymmetric apoptotic patterns associated with local hyperconfluent regions. Further, groups of aligned elongated cells were observed spanning the 200 μm-diameter collagen islands which are inconsistent with the predicted stress fields and have not been reported for epithelial cells or cell lines cultured on similar patterned protein islands.

The goal of this work is to uncover the mechanisms underlying the observed heterogeneous collective cell behaviors within micropatterned VIC aggregates. Circular stamps are employed to provide a uniform radially symmetric global constraint, and relatively large 400 μm-diameter collagen islands were chosen to minimize the spanning of single cells across the patterns. We use long-term time-lapse imaging to follow the evolution of individual aggregates over four days as substantial heterogeneity between aggregates has been observed even in a single dish of uniformly printed cell islands. Short-term time-lapse is used to observe cell-cell interactions in real time and to quantify cell velocity and traction stresses in distinct sub-regions over the span of hours. To determine if cells act independently or collectively in response to an imposed global stimulus, we cyclically stretch aggregates and quantify changes in orientation and elongation. As Yes-associated protein (YAP) is implicated in mechanosensing in many cell types (Aragona et al. 2013, Calvo et al. 2013, Codelia et al. 2014, Das et al. 2016), we investigate the relationship between YAP activation and apoptosis stemming from the collective cell behavior. The knowledge obtained from this study provides further insight into how collective cell behaviors drive biological phenomena that are implicated in many developmental and pathological conditions.

## 2. Methods

### 2.1 Substrate preparation

Microcontact printed 400 μm diameter circular protein islands were formed by coating plasma-treated and 70% EtOH pre-rinsed polydimethylsiloxane (PDMS) stamps with collagen and placing them onto untreated 22x22 mm square coverslips for 1 hr; a 50 g weight was lightly applied to create uniform pressure on the stamp (Fig. S.1). The collagen solution consisted of 25 μL of 4 mg/mL collagen, 75 μL of 0.1 M acetic acid, 900 μL sodium acetate buffer, and sodium periodate. Prior to transfer, excess collagen was removed from the stamp using a combination of air drying and nitrogen stream. In a subset of experiments, the uniformity of the circular collagen prints was verified with the addition of 1 μL Alexa Fluor-488 carboxylic acid, succinimidyl ester (A20000, Invitrogen) to the collagen solution for 1 hr at room temperature. To determine the effects of island size, we used 200 μm and 600 μm circular patterns in a subset of experiments.

Circular collagen patterns were printed onto polyacrylamide (PA) gels by indirect microcontact printing (Cirka et al. 2017). Briefly, for each substrate, 50 μL of PA solution (acrylamide:bisacrylamide (Biorad) of percentages of 7.5/0.24 for ∼20 kPa) were pipetted onto an activated coverslip. The coverslips were activated by soaking in 1.5% (aminopropyl) trimethoxysilane solution for 30 min then in 0.5% glutaraldehyde solution overnight at 4 °C and dried completely before use. After placement of PA solution on activated coverslips an inactivated collagen microcontact printed coverslip was gently placed on top. Following 12 min of polymerization of the PA-gel, the coverslips were separated by a razor blade. A modulus of ∼20 kPa was chosen because it is in the range of reported stiffness values for healthy and diseased valves (Kloxin et al. 2010, Wang et al. 2012).

### 2.2 Traction force microscopy

To quantify the shear stresses that the cells apply to the surface of the substrate via traction force microscopy (TFM), 0.2 μm red fluorescent micro-beads were coated on plasma treated glass coverslips, allowed to dry, and then applied to the top surface of the PA gel solution during polymerization. Collagen patterns were then stamped onto the PA gel using direct microcontact printing methods as previously described (Cirka et al. 2016). At various timepoints, phase and fluorescent images were acquired to determine the aggregate borders and bead locations, respectively. Cells were then trypsinized, and a reference image of the bead locations was acquired. Displacements of the beads were calculated with a custom MATLAB code and input into a finite element model (modulus: 19 kPa; Poisson ratio: 0.4; material property: linear elastic) to calculate the stresses on the surface of the gel (ANSYS Inc.) (Cirka et al. 2016).

### 2.3 Dynamic stretching

To determine the effect of cyclic stretching on collective behavior in aggregates, PA gels were affixed in each well of a 16-well compliant Elastosil culture plate (CellScale), patterned using indirect microcontact printing, and seeded with cells. After 24 hr post-seeding, the substrates were stretched 10% uniaxially at 1 Hz for 8 hr using a MechanoCulture FX (CellScale). PA gels were attached to the compoliant substrate by treating wells with 1.5% (aminopropyl) trimethoxysilane solution for 5 min, drying, and then treating with 0.5% glutaraldehyde solution for 5 min. After removing the liquid, the wells were dried with nitrogen stream, 4 μL of PA were placed at the bottom of the well and covered with a collagen patterned circular 5 mm coverslip. The wells were then transferred to a vacuum chamber filled with nitrogen for 45 min to facilitate polymerization, then diH_2_O was added to the wells for 30 min to promote detachment of the coverslips from the PA gel surface.

### 2.4 Cell culture and media

Aortic VICs were isolated from porcine hearts obtained from a local abattoir (Blood Farm, Groton, MA) using previous protocols (Gould et al. 2010). Porcine VICs are primary fibroblastic cells and were chosen as VIC aggregation is implicated in the pathology of CAVD, in addition to their similarity to human VICs. VICs at passages 3-6 were seeded at 12,500 cells/cm^2^ (Cirka et al. 2017). In one set of experiments, neonatal human dermal fibroblasts (courtesy G. Pins, Worcester Polytechnic Institute) were used with the same passage range, cell density, and culture conditions as VICs. The cells were cultured in DMEM supplemented with 10% FBS and 1% Antibiotic/Antifungal at 37 °C with 10% CO_2_. Media were exchanged every 48 hr. In cell-cell interaction inhibition experiments, calcium was depleted from DMEM with 1 mM ethylenediaminetetraacetic acid (EDTA), a concentrated reported to not affect cell viability (Voccoli et al. 2014).

### 2.5 Stable YAP-VIC cell line

To determine if apoptosis is YAP-dependent, YAP-6A, a constitutively active version of YAP that cannot be inhibited by LATS, was obtained from Addgene (#42562), packaged into lentivirus particles, and transduced into VIC cells. Lentiviral particles were generated by transfecting 293T cells with pLX304-YAP-S6A-V5, pSPAX.2, and PMD2.G. After 48 hr, the supernatant was collected and filtered through a 45 μm sterile filter. Viral supernatant was then mixed 1:1 with culture media, added with 1 μg/mL of polybrene (Millipore), and incubated with VICs overnight. The next day, cells were fed with culture media, and after 24 hr, cells were selected with 1 μg/mL of Puromycin until all control (uninfected) cells died after approximately 48 hr. Expression of the YAP S6A-V5 construct was visualized in pooled puromycin-resistant cells through immunofluorescence using anti-V5 (Invitrogen).

### 2.6 Imaging and immunohistochemistry

Individual aggregates were tracked for four days utilizing a motorized stage and automated position tracking software (Zeiss Axiovision 4.8.2 SP1) on an inverted microscope (Axiovert 200M, Zeiss) and imaged with phase contrast every 24 hr at 10x (AxioCam MRm camera; 1.4 MP, 1388 X 1040 pixels). To track individual aggregates at specific locations, coordinates of each aggregate were saved with respect to a reference point at the top left corner of the coverslip on each day. Samples were incubated for 30 min in standard media with CellEvent Caspase 3/7 (1:400, C10423, Invitrogen) prior to imaging to observe apoptosis.

YAP staining was completed following published methods (Ma et al. 2017, Dutta et al. 2018). VIC aggregates were fixed in 4% paraformaldehyde for 45 min at room temperature, rinsed twice in PBS, and permeabilized using 0.1% TritonX-100 in PBS for 1 hr. Samples were then blocked with 5% bovine serum albumin (BSA) overnight at 4°C to minimize nonspecific protein binding. Samples were stained with Anti-YAP in 5% BSA (1:250, mouse, sc-398182, Santa Cruz) primary antibody and incubated for 1 hr at room temperature. Samples were rinsed in PBST (0.5 wt% Tween-20 in PBS) two times for 10 min following incubation with primary antibody. Samples were then incubated in secondary antibody with goat anti-mouse AlexaFluor 647 (1:200, A21241, Invitrogen) while counterstained with Alexa Fluor 488 Phalloidin (1:100, A12379, Life Technologies) to visualize F-actin and Hoechst 33342 (1:200, H3570, Life Technologies) to visualize nuclei in 1% BSA for 1 hr at room temperature. Samples were rinsed three times for 10 min with PBST, mounted, and imaged using a BioTek Cytation Gen5 inverted microscope. To quantify YAP activation, the colocalization of YAP and Hoechst signals was quantified as nuclear, nuclear/cytoplasmic, or cytoplasmic in the hyperconfluent bands identified in phase images.

To visualize migration of individual cells within the aggregates, 20% of cells were treated with CellTracker™ Green CMFDA Dye (5:1000, C2925, Invitrogen) and imaged via fluorescence microscopy (Axiovert 200M, Zeiss). Only staining a portion of the cells is required so that individual cells can be tracked when cells become dense. To visualize if the collagen island remained after aggregates detached, a subset of samples was fixed and stained with picrosirius red/fast green (120M1516V, Sigma-Aldrich) and imaged with transmitted light and color camera (Cytation 5, BioTek).

To quantify proliferation, Click-iT™ Plus EdU Cell Proliferation Kit for Imaging, Alexa Fluor™ 488 dye was used per manufacturer’s instructions (C10637, Invitrogen). We performed both a 24 hr-long exposure to visualize proliferation over an entire population doubling and a 1-hr pulse experiment for short-term visualization with minimal migration away from the site of proliferation.

### 2.7 Quantification of confluency level, cell density, and cell alignment

To quantify the observed confluency levels in each aggregate, the area of the cells in phase images was measured using ImageJ and divided by the total area of the collagen island in the same image. To quantify the average density of cells in different confluency levels, the number of the cells in each aggregate was divided by its area.

To quantify the alignment of cells in live cultures before and after stretch, F-actin was stained using CellMask™ Actin Tracking at 1:1000 dilution (A57249, Invitrogen). As this staining of live cells is not as efficient as phalloidin staining of fixed and permeabilized cells, only a portion of the cells are stained brightly. Further, imaging through the 400 μm-thick silicone well and the∼200 μm-thick PA gel degrades the quality of the images. Together, these limitations make quantification of the stress fiber angular distribution problematic; however, as individual cells could be identified clearly, we were able to track these individual cells before and after stretch and quantify their direction and elongation by tracing each stained cell and calculating the angle and elongation of the stained fibers with the Directionality tool in ImageJ.

### 2.8 Statistical Analysis

Data are presented as mean and standard deviation. One-way ANOVA was performed to evaluate the significance of cell density values between the five different confluency levels. Holm-Sidak post-hoc test was used with all pairwise comparison. Student’s t-test was used to evaluate the significance in differences of cell speed between confluent and hyperconfluent regions. Paired t-tests were used to evaluate of the significance of changes in the cell orientation and aspect ratio before and after stretch. Analyses were completed in SPSS, and a p-value of less than 0.05 was considered statistically significant.

## 3. Results

### 3.1 Aggregates have variable rates of progression towards confluency

Across the span of 4 days, 53 aggregates were tracked following initial seeding. On the 400 μm-diameter collagen islands containing 50-300 cells, cells migrate, proliferate, and fill the islands at varying rates. As the density of cells was variable within each aggregate, we manually categorized aggregates in terms of confluency level and measured the percentage of the island area that is covered with cells (Fig. 1A). In the “sparse” stage, relatively large regions of islands are visible between cells and coverage is 80.0±9.0%. In the “pre-confluent” stage, cells fill most of the circular area (89.6±5.1% coverage) and only small areas of the surface are visible. At the “confluent” stage, cells cover almost the entire area (95.6±1.7% coverage), and cell density is approximately uniform. The “hyperconfluent” stage is characterized by regions in which cells are highly dense and rounded, although there are still small gaps in some aggregates (97.3±0.8% coverage). In the “self-detached” stage, highly dense areas are formed when portions of the aggregate pull away from the culture surface. The average cell density correlates with the confluency level (Supplemental Fig. S.2A), although there is not a one-to-one correspondence due to the heterogeneity within each aggregate e.g., some aggregates with highly dense regions are categorized as hyperconfluent yet have an average cell density in the confluent range.

**Figure. 1.**
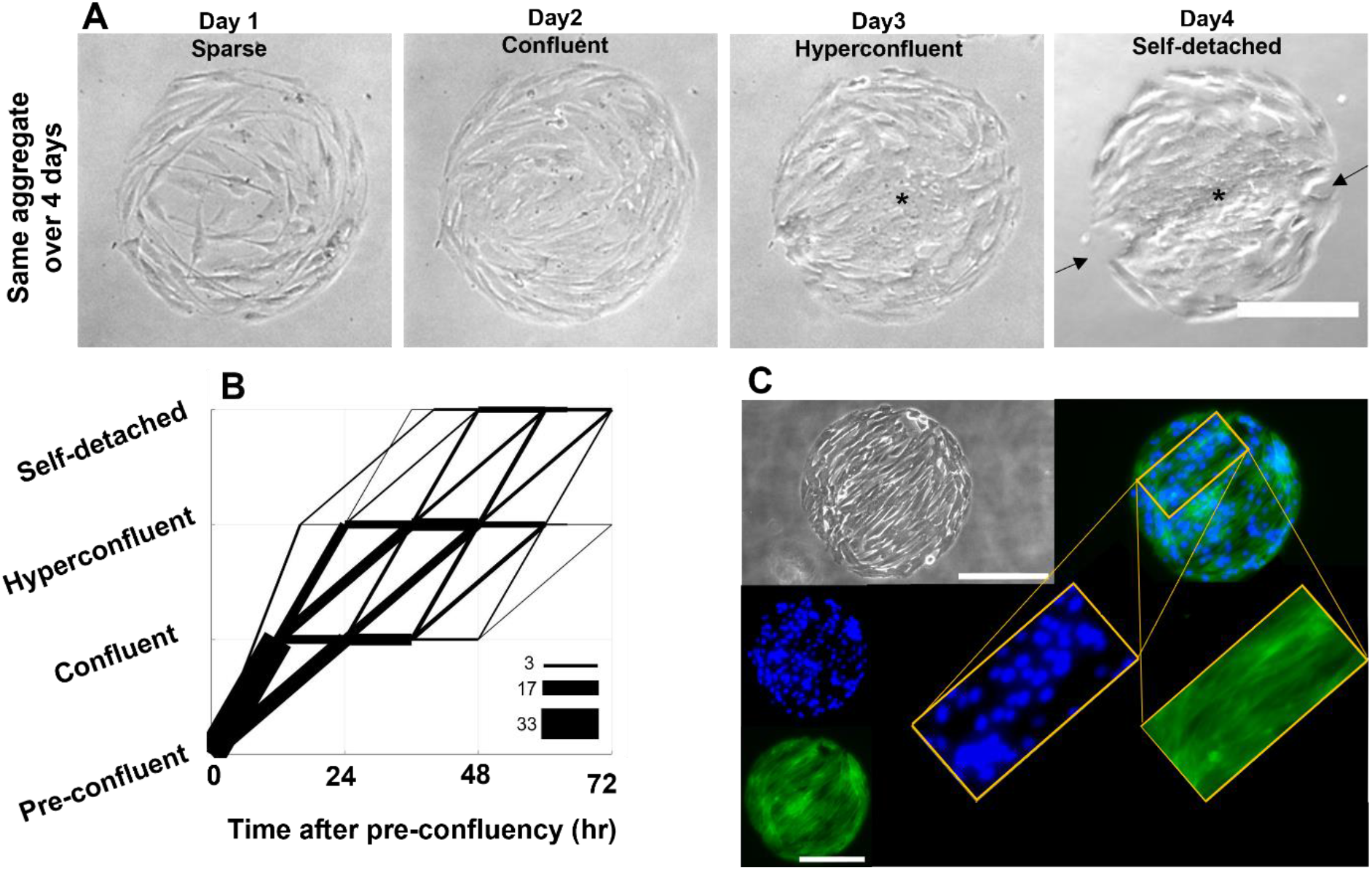
The progression between levels of confluency varies in time from aggregate to aggregate. A) Time-lapse of the same aggregate over 4 days showing the evolution of its confluency starting sparce 24 hr after seeding, proceeding to confluent at day 2, hyperconfluent on day 3 (* indicates high density region), and finally self-detachment on day 4 (arrow indicate regions detached from substrate). B) Plot of levels of confluency versus time adjusted to time when each particular aggregate reached the pre-confluent level to remove variability in initial seeding in a single dish (n = 53 aggregates; thickness of lines indicates number of aggregates following a given path). The majority of aggregates take 12 hr to reach confluency, yet there remains high variability in confluency rates from pre-confluent to higher levels of confluency after removal of heterogeneity in initial seeding (see other examples in Fig. 2 and Supplemental Fig. S.2). C) Local clustering of aligned cells forms banding and hyperconfluent regions. F-actin staining (green) shows the direction of the band, and high density of nuclei (blue) demonstrates hyperconfluency. The phase image in panel C (gray) was captured after fixation and permeabilization which creates spaces between cells not seen in live imaging in panel A. Scale bar = 200 μm.

To account for potential inconsistencies in initial seeding, we set the pre-confluent level to be “time zero.” Despite this data shift, we still observe substantial variability in how long it takes aggregates to progress through confluency levels (Fig 1B). The majority of aggregates progress from pre-confluent to confluent within 12 hr (as indicated by the thickness of the line). Note that the aggregate shown in Fig. 1A corresponds to a line in Fig. 1B where every 24 hours the level of confluency moves one higher. Rather than seeding density, it appears that banding plays a predominate role in variability in timing. When a band is initiated in the pre-confluent stage, aggregates progress quickly to hyperconfluence, whereas if cells do not form bands until later phases, progression is slower (Supplemental Fig. S.2B, Fig. S.2C, respectively). These multicellular bands can be observed more clearly in phase imaging following fixation when individual cells separate due to the dehydration process, and fluorescent staining shows continuous F-actin structures spanning regions with high numbers of nuclei (Fig. 1C).

### 3.2 Local hyperconfluency occurs following band formation

Despite the simple circular shape of the collagen islands, the patterns in relatively few aggregates were radially symmetric with circumferential alignment at edges and isotropic in the center (Fig. 2A). Rather, in most aggregates, multicellular banding was observed disrupting this radial symmetry, even at early timepoints (Fig. 2B left panels). After 4 days, pronounced hyperconfluent bands were observed in all aggregates (Fig. 2B right panels). If banding is pronounced at the pre-confluent stage (Fig. 2C left panels), the band becomes highly dense within 24 hr and leads to self-detachment of cells from the substrate at the two endpoints of the banded region (Fig. 2C right panels).

**Figure. 2.**
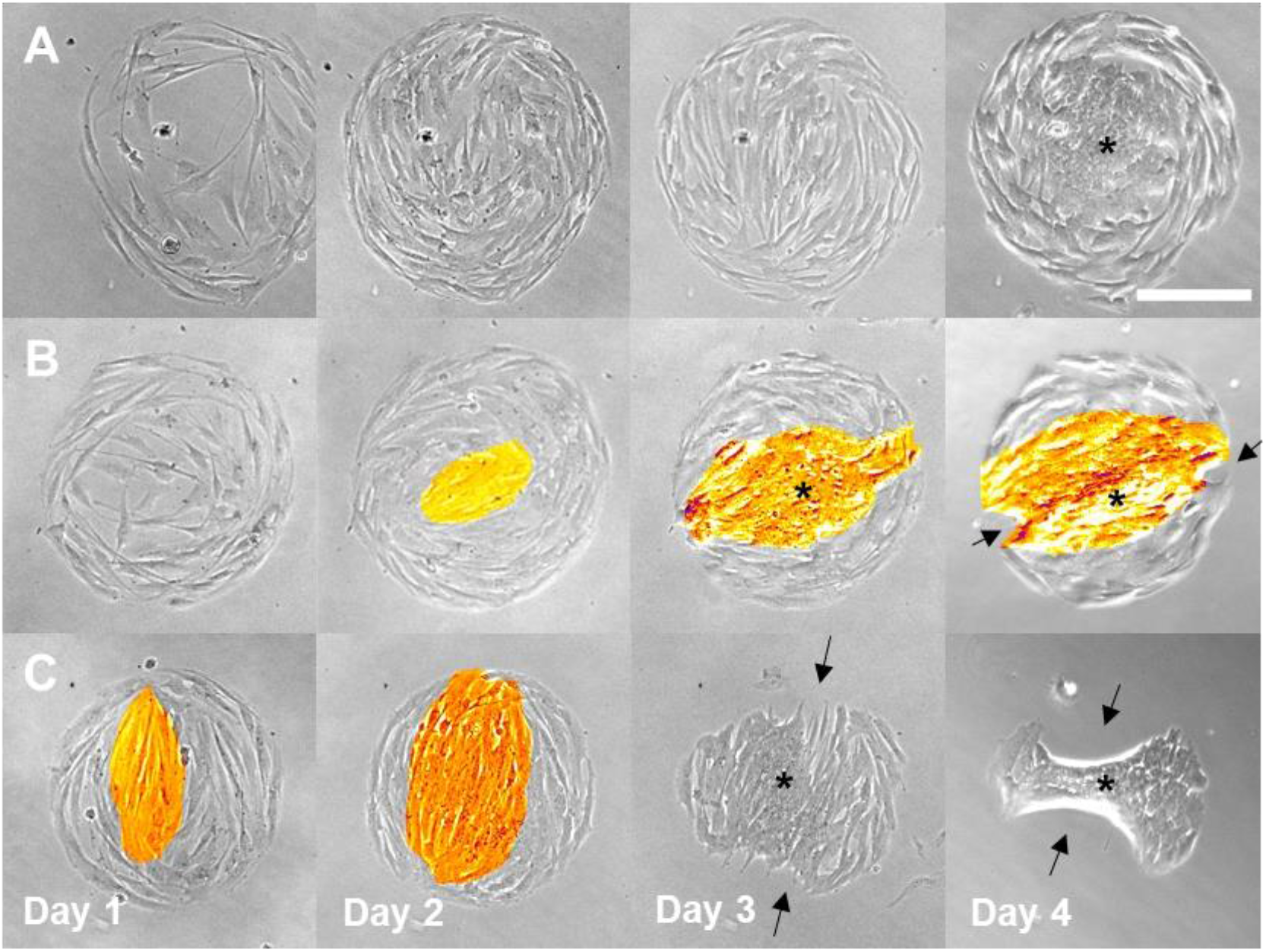
Three examples of different time courses of confluency progression in aggregates. A) Aggregate with relatively uniform cell distribution exhibiting circumferentially aligned cells around the edge and high density in the center by day 4 but no banding or self-detachment. B) Aggregate with formation of band starting day 2 and leading hyperconfluency by day 4 and slight self-detachment by day 4. C) Aggregate with banding occurring early which proceeds quickly to hyperconfluency and self-detachment by day 3. Banding of aligned cells is highlighted in orange color, * indicates high density region, and arrows indicate regions detached from substrate. Scale bar = 200 μm.

In the majority of aggregates, bands form a straight line from one edge of the aggregate to the other. Less often the bands are curved, only span a small internal region, or connect into triangular patterns (Supplemental Fig. S.3). Most bands eventually lead to some degree of self-detachment within the 4-day culture period.

### 3.3 Banding occurs regardless of aggregate size

In previous studies, we microcontact-printed 200 μm-diameter protein islands and observed VICs forming confluent aggregates with cells generally oriented circumferential at the edges but highly heterogeneous orientation and cell area in the central region (Goldblatt et al. 2020). In this study, we used 400 μm-diameter to minimize the “edge effect,” yet we still observed highly heterogenous structures within most aggregates. To further lessen the proximity to the edges, 600 μm-diameter islands were created, and similar hyperconfluent bands formed in these large aggregates as well (Fig. 3). This finding, in conjunction with our time-lapse studies where cells aligned with individual elongated cells in aggregate centers, indicates that bands are not forming by cells interacting with the edges. To determine if this behavior is unique for VICs, we cultured primary human dermal fibroblasts on 400 um islands, and we observed similar banding and hyperconfluency (Supplemental Fig. S.4).

**Figure 3.**
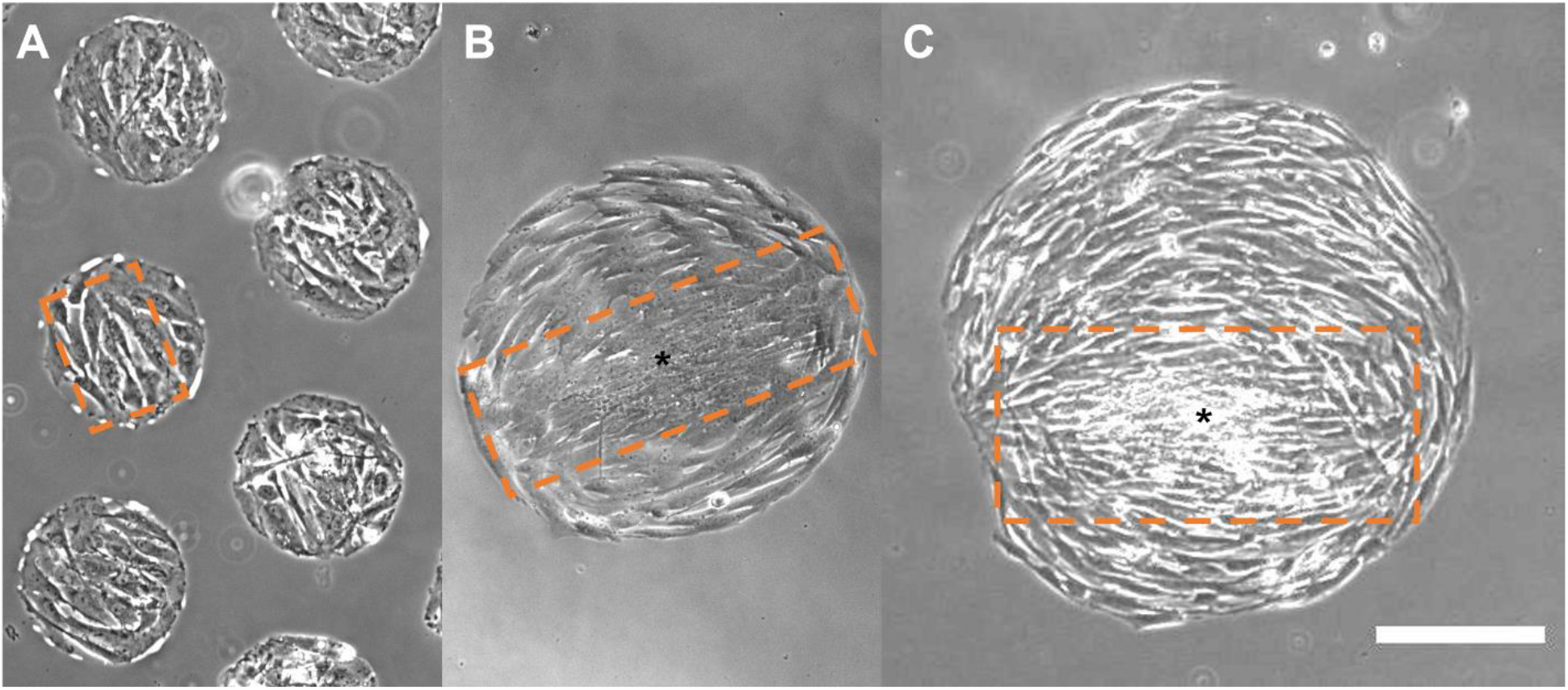
Banding and hyperconfluency occurs independent of aggregate size as observed in A) 200 μm, B) 400 μm, and C) 600 μm day 4 aggregates. Banding is highlighted by dashed orange rectangle and * indicates high density region. Scale bar = 200 μm.

### 3.4 Cells migrate to connect with bands and then move in coordination

To better understand how cell migration plays a role in band formation, we performed 4.5 hr time-lapse imaging of aggregates with approximately 20% of cells fluorescently labeled so that individual cells could be tracked. Following 162 cells, we observed that cells in relatively sparse regions moved independently, whereas cells within bands moved coherently. The vast majority of cells (∼96%) remain within their respective regions (sparse, band, or edge) over the 4.5 hours (Supplemental Movie M.1). Five cells in sparse regions merged with bands and one cell joined a band from the edge. Four cells from relatively sparse regions joined the ring of aligned cells along the edge in the absence of any band. Although the behavior of cells in different regions were qualitatively different with cells in sparse regions migrating relatively independently and the cells in hyperconfluent regions oscillating in sync, the average speed in less dense regions (0.14 ± 0.01 μm/min) was not significantly different than in hyperconfluent regions (0.12 ± 0.02 μm/min, t-test, p = 0.07).

To understand the events leading to band formation, we imaged aggregates every 30 minutes over 24 hours. Cells under lower confluency levels display a fluid-like behavior with edge cells rotating around the circumference and internal cells beginning to align to one another and forming bands (Supplemental Movie M.2A). As confluency progresses, motility within aggregates decreases and cell migration is relatively uncoordinated (Supplemental Movie M.2B). Finally, during hyperconfluent stages, cells move back and forth together with limited migration similar to an oscillation (Supplemental Movie M.2C). Cell proliferation is observed in the phase images at all confluency levels with the exception of hyperconfluent regions. Propidium iodide staining at the beginning of time-lapse experiments suggests that a small amount of apoptosis occurs throughout sub-confluent and confluent aggregates, whereas the rate of cell death is much higher in the middle of hyperconfluent bands. The propidium iodide signal dissipates with time due to photobleaching so is only seen at the early timepoints.

### 3.5 Local hyperconfluency is inhibited by depleting calcium

To test whether band formation is driven by intercellular force transmission, we decreased cell-cell adhesion by precipitating calcium from the cell culture medium. After four days in culture, the last two of which were in calcium-free media, hyperconfluent regions did not form, and we did not observe any self-detachment (Fig. 4). However, local regions of cell alignment were still observed. This local alignment may have been established in the first two days of culture in standard medium which is necessary for forming aggregates as proliferation is calcium dependent (Kahl et al. 2003).

**Figure 4.**
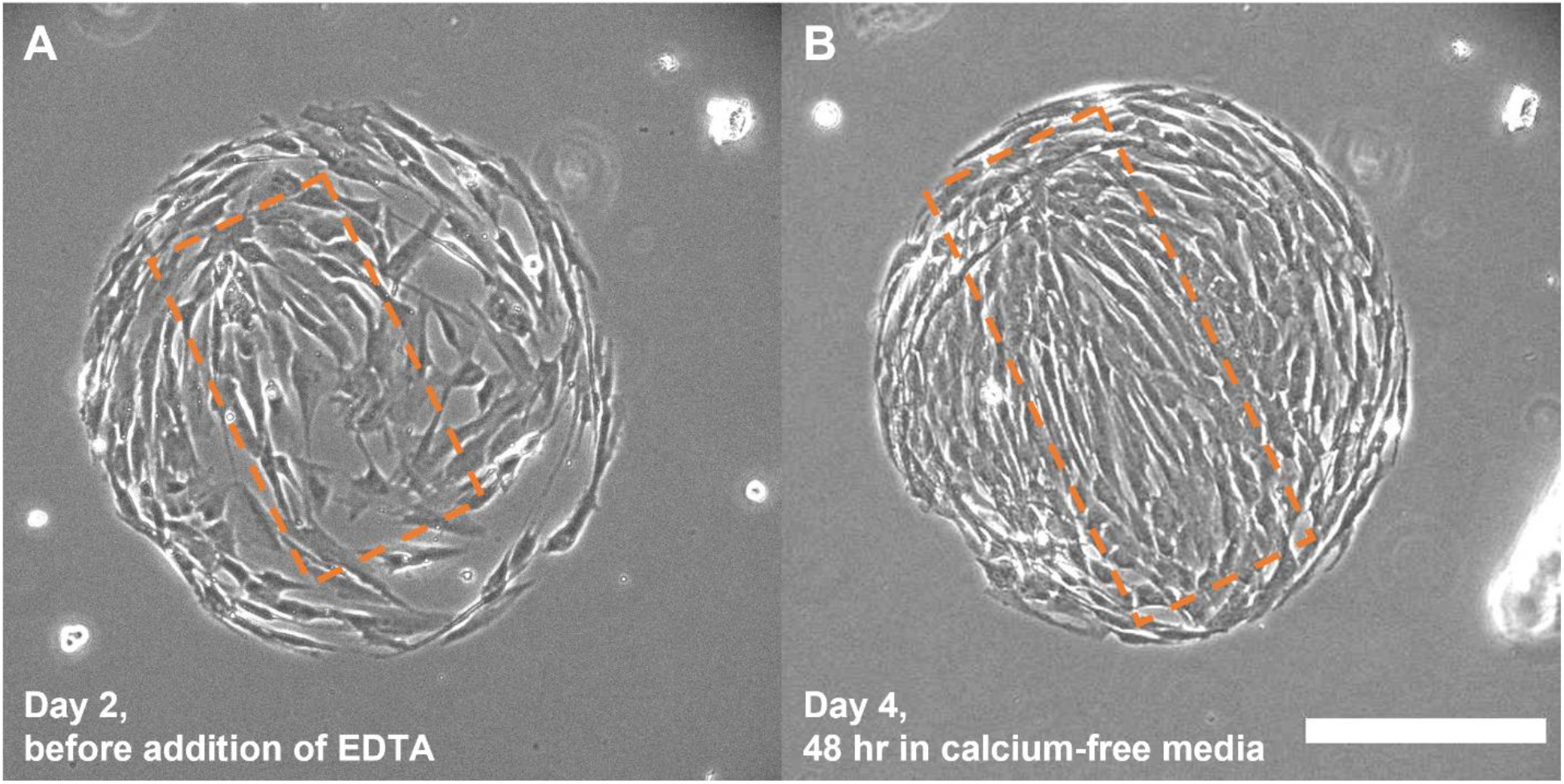
Inhibiting cell-cell adhesion via calcium depletion prevents hyperconfluency, although banding is still observed. A) A day 2 aggregate before addition of EDTA shows pre-alignment of the cells. B) The same aggregate on day 4 after 2 days in calcium-free medium shows aligned cells in a band; however, the cells are not tightly contacting and hyperconfluent regions are absent. Banding is highlighted by dashed orange rectangle. Scale bar = 200 μm.

### 3.6 Hyperconfluent bands increase local traction stresses

Unlike traction stress maps for homogeneous aggregates in which the traction is high and relatively uniform at the edges (Li et al. 2009), we observe that the bands accentuate the traction stresses at their endpoints (Fig. 5). When aggregates are confluent, traction “hot spots” are relatively low (Fig. 5A, B) compared to when large hyperconfluent bands form (Fig. 5C, D). Analysis of the principal stress vector directions indicate that stresses are highly anisotropic and predominantly in the direction of bands (Fig. 5 insets 1-4). Full vector fields are provided in the Supplemental Information (Supplemental Fig. S.5). Time-lapse traction stress maps show that stresses dynamically change in magnitude and location over 5 hr with cell movement, but peak traction stresses remain at the band ends (Supplemental Movie M.1). Eventually, high traction stresses at band ends lead to self-detachment by exceeding the strength of the collagen-PA substrate bond/entanglement (Supplemental Fig. S.6).

**Figure 5.**
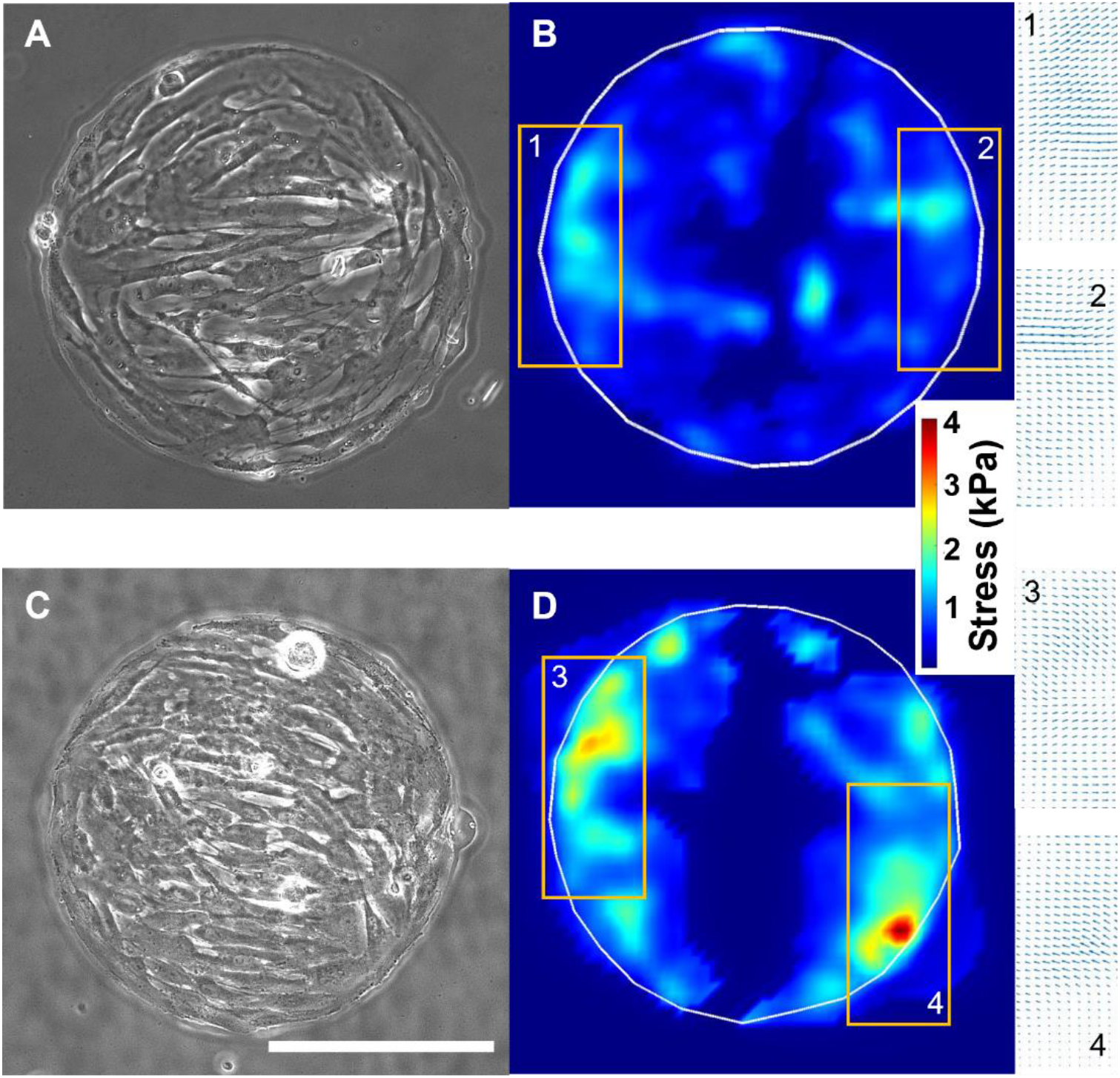
Heterogeneous traction stress fields are produced by alignment and banding of cells and peak stresses increase with cell density. A) Phase image of a sparse aggregate shows initial alignment of cells. B) TFM analysis of the sparse aggregate shows low magnitude stress hotspots at ends of aligned cells with inward direction of applied stress (insets 1 & 2). C) Phase image of a confluent aggregate shows strong banding. D) TFM analysis of confluent aggregate shows high magnitude stress hotspots with inward direction of applied stress (insets 3 & 4). Scale bar = 200 μm.

### 3.7 Cytosolic YAP indicates low tension in hyperconfluent bands

Despite measurement of high traction stress at the ends of the bands, we hypothesized that the force per cell is low within the bands due to the large number of cells acting in parallel in these hyperconfluent regions. As an indirect measure of cell stress, we stained for YAP, a mechanosensitive protein which is shuttled preferentially to the nucleus under high tension and remains cytosolic in low tension environments (Aragona et al. 2013). In our aggregates, cells inside of bands have more cytosolic YAP (Fig. 6B, inset 1) compared to cells outside of bands, which have more nuclear YAP (Fig. 6B, inset 2). At band endpoints, where cells are more elongated, YAP is predominantly nuclear (Fig. 6B, inset 3). We did not observe any purely cytosolic YAP.

**Figure. 6.**
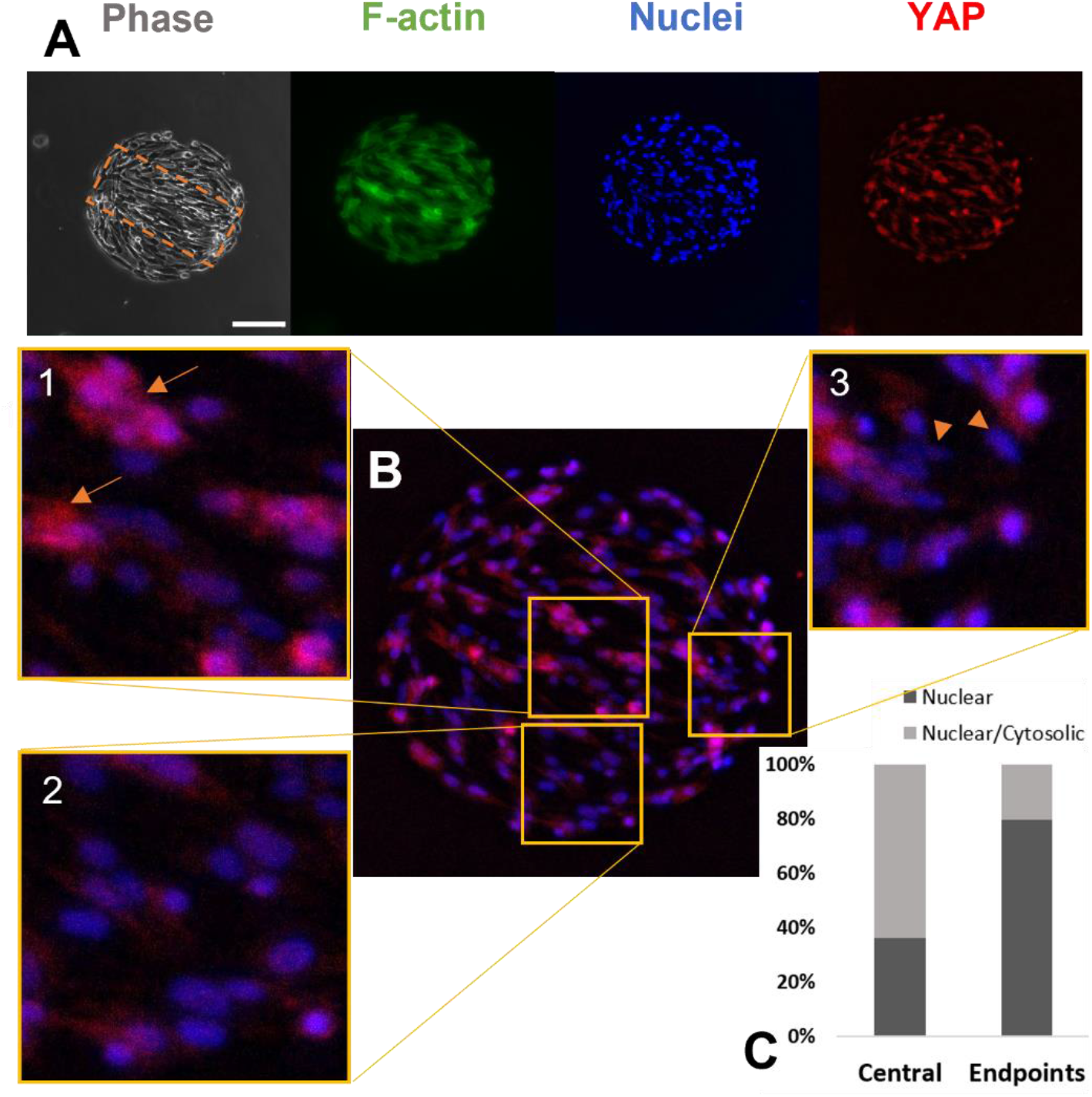
A greater proportion of cytosolic YAP is observed in the center of hyperconfluent bands than at the ends. A) Phase image of fixed and permeabilized and fluorescent images of F-actin, nuclei, and YAP in a hyperconfluent aggregate exhibiting banding (dashed orange box). B) YAP/Hoechst merged image shows mixed cytosolic/nuclear YAP in the hyperconfluent center (inset 1), outside of band (inset 2), and nuclear YAP at the endpoint of the band (inset 3). Arrows point to examples of cytosolic/nuclear YAP, and arrowheads point to examples of nuclear YAP. Scale bar = 200 μm. C) Quantification of nuclear (N) and nuclear/cytosolic (N/C) YAP in the center and end of bands (n = 212 cells, 4 aggregates); there was no purely cytosolic YAP observed.

### 3.8 Cells in bands reorient towards stretch but not completely

To examine the response of cells to external loading, we applied 10% cyclic uniaxial stretch for eight hours at 1 Hz frequency. Due to the heterogeneity of band locations, we tracked 38 individual aggregates before and after stretch rather than averaging overall cell behavior between aggregates. Actin fibers were with a live stain taken up by cells and imaged in compliant stretch wells without fixation; thus, since not all cells were not stained equally, we manually tracked and quantified the morphology of 56 individual cells within those aggregates before and after stretch.

In general, the cells reorient towards the direction of stretch but are not able to completely reorient to the stretch axis, and the extent of cell reorientation depends upon initial angle of the bands. For cells inside bands originally aligned within ±15° of the direction of stretch (Fig. 7A, circled cells), the alignment of the cytoskeleton becomes more pronounced, and the cells reorient slightly from 5.2°±7.9° to 2.7°±3.2° (p=0.45, n=9). The aspect ratio of these cells in the direction of stretch increases significantly from 3.7±1.4 to 6.3±3.2 (p=0.04). When bands are at a moderate angle (15° to 45°) relative to the stretch direction, cells reorient significantly towards the direction of stretch but not fully, from 28.0°±13.2° to 18.4°±13.8° (p=0.001, n=23), and aspect ratio trends higher from 2.5+-0.7 to 3.0+-1.3, but the change is not significant (p = 0.53) (Fig. 7B, Supplemental Fig. S.7A). When bands are at a large angle relative to the stretch direction, initially at 45° to 90°, a subset of cells reorient towards the stretch direction significantly from 59.1°± 8.8° to 30.0°±18.3° (p=0.003, n=9), whereas in others the cytoskeleton is disrupted, and the cells appear more rounded and the orientation was not quantified (Fig. 7C, Supplemental Fig. S.7B).

**Figure 7.**
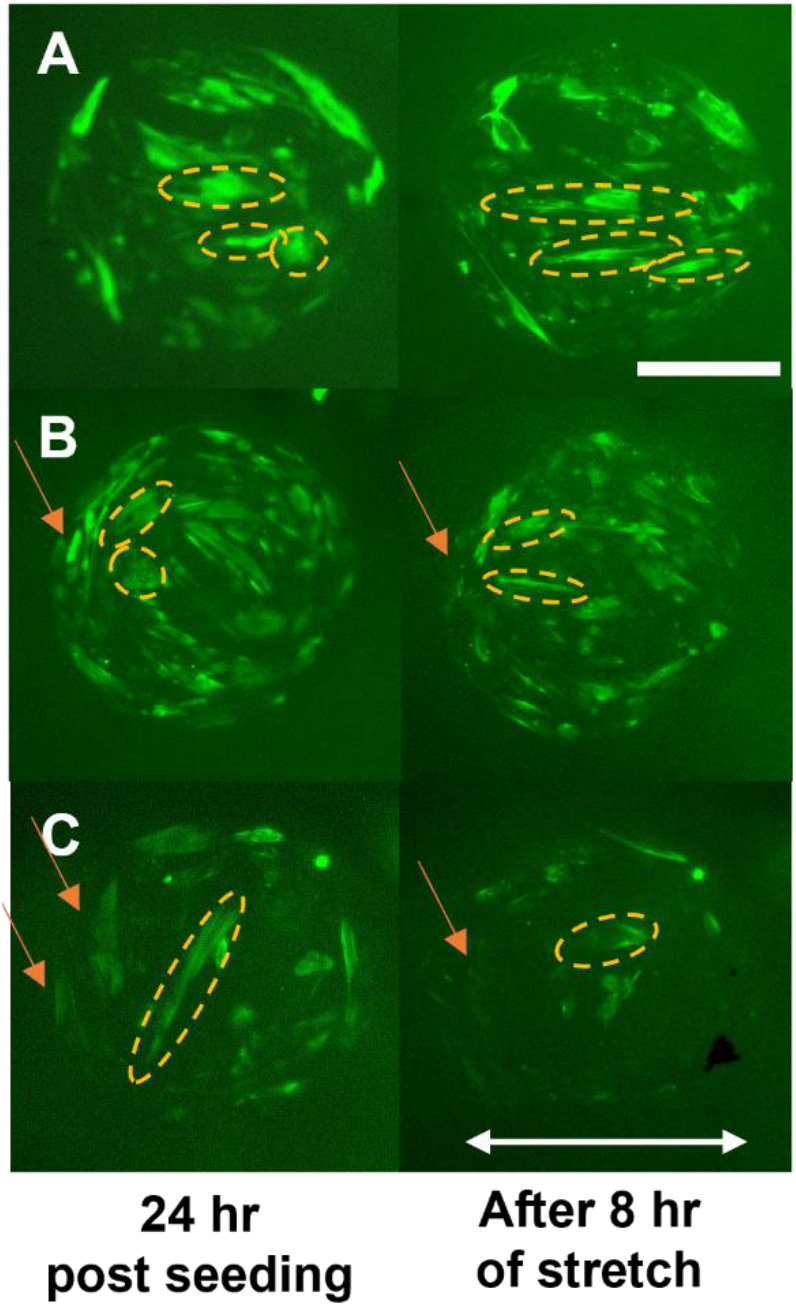
Dynamic stretch reinforces F-actin alignment for cells in the direction of stretch and disrupts the cytoskeleton of cells aligned away from the stretch axis. A) Representative example of cells within a band roughly in the direction of stretch where the F-actin is intensified, and the cells are elongated after stretch. B) Representative example of cells within bands that are oriented within 45° of the stretch axis showing reorientation towards the direction of stretch. C) Representative example of a highly elongated cell within a band oriented greater than 45° from the stretch axis becoming more rounded and reorienting towards the stretch direction. Panels B and C also show that cells aligned along the edge perpendicular to the stretch direction retract (orange arrows). For all panels, F-actin of live VICs was stained with CellMask™ Actin before and after application of 10% uniaxial stretch for 8 hr at 1 Hz; brightly stained cells inside of bands are outlined by ellipses to show change in orientation and elongation. Scale bar = 200 μm.

Cells on island edges that align with the global circular constraint and are in the direction of stretch (top and bottom of the images) are relatively stable and reorient only slightly from – 2.5°±16.0° to 1.8°±12.0° (p=0.30, n=16). Conversely, for cells that align to the edges that are perpendicular to the stretch (left and right sides of the images), the cytoskeletons are disrupted thus the reorientation of the perpendicular cells could not be quantified (Fig. 7B, 7C); this cytoskeletal disruption was observed in 20 of the 24 tracked aggregates where perpendicular cells were observed at the edges at pre-stretch.

### 3.9 Proliferation occurs throughout aggregates except in hyperconfluent bands

In contrast to studies showing proliferation of cells predominantly at the edges of constrained epithelial cell monolayers (Li et al. 2006, Li et al. 2009, Aragona et al. 2013, Silver et al. 2020), we observe proliferation throughout our confluent aggregates. Images from 1 hr EdU pulse experiments show no distinct pattern of proliferation (Fig. 8A) even when images of positive cells from multiple aggregates are binarized and stacked (Fig. 8B). The 24 hr EdU exposure experiments indicate that proliferation does not occur in hyperconfluent bands (Fig. 8C, 8D).

**Figure 8.**
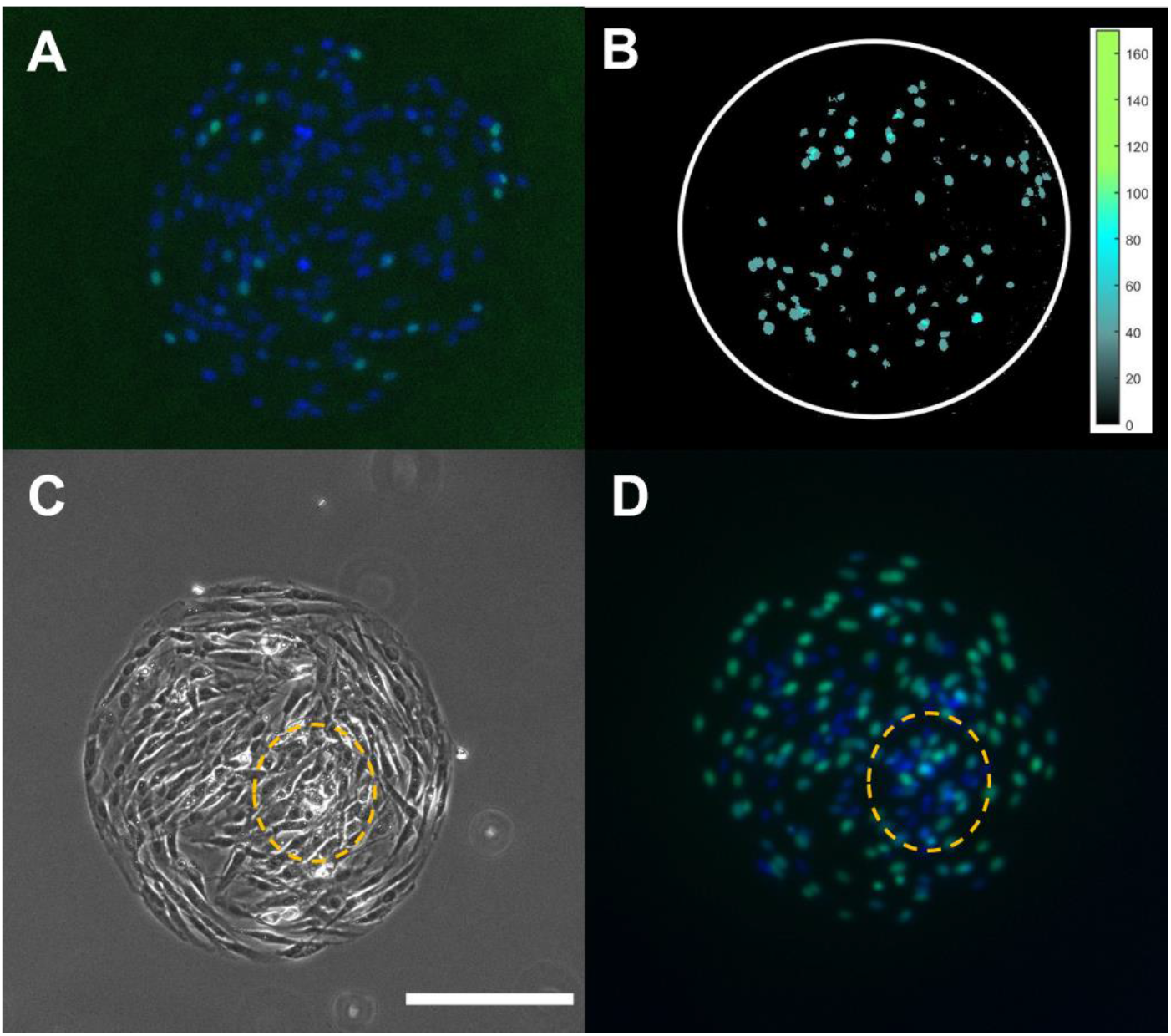
Proliferation occurs throughout confluent aggregates except in high density regions. A) Representative aggregate after 1 hr pulse staining for proliferation with EdU. B) Overlay of binarized images of proliferation for 4 different aggregates after 1 hr EdU pulse shows proliferation throughout confluent aggregates. C) Representative phase image of a fixed and permeabilized aggregate with a hyperconfluent region (orange outline). D) After 24 hr EdU exposure, proliferation occurs throughout the aggregate except in the hyperconfluent region. Fluorescent images are merged Hoechst (blue) and EdU (green) images. Scale bar = 200 μm.

### 3.10 Apoptosis increases in high density regions in a YAP-dependent manner

As reported above, in time-lapse videos with PI staining (Supplemental Movies M.2A, B, C), we observe a low incidence of cell death throughout aggregates except within hyperconfluent regions. We confirmed that this cell death is likely due to apoptosis by staining for cleaved caspase 3/7. Within the first 24 hr after pre-confluency, approximately 20% of aggregates show a caspase positive signal indicating apoptosis. Most caspase-positive aggregates are confluent aggregates as few hyperconfluent aggregates are present at this time (Fig. 9). At 24-48 hr after pre-confluency, almost half of the aggregates show caspase activity, and most of these aggregates are hyperconfluent. At 48-72 hr after pre-confluency, nearly all aggregates show a positive caspase signal with the majority having detached from the surface.

**Figure 9.**
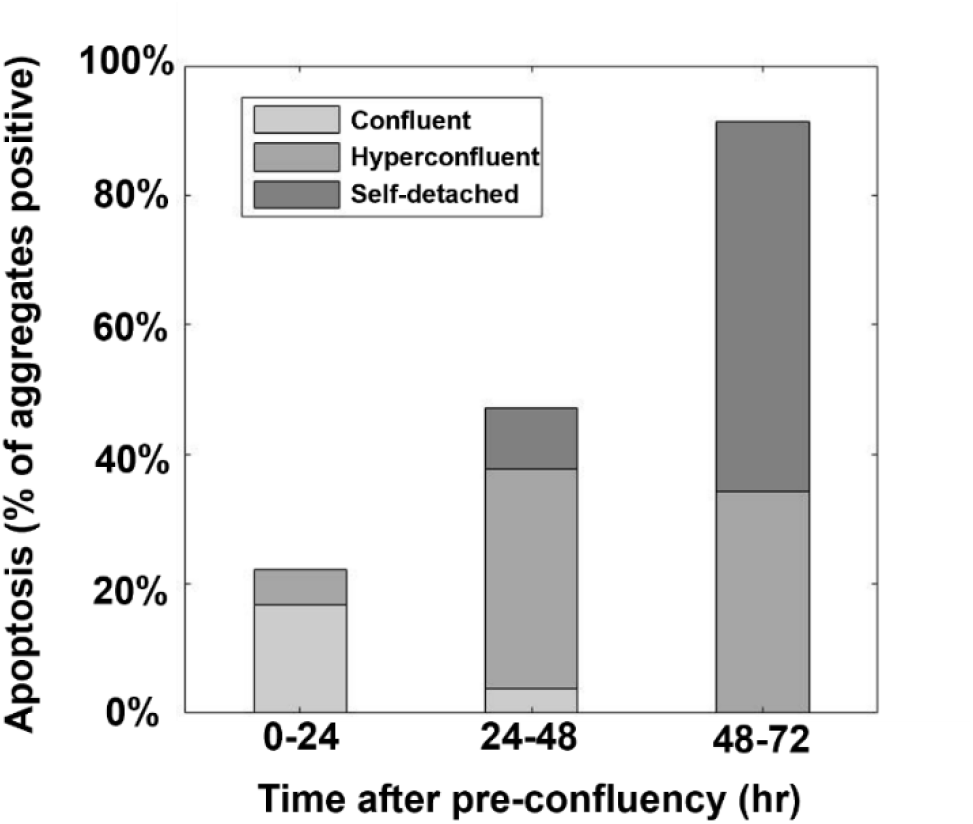
Apoptosis increases with culture time. A small proportion of confluent aggregates have apoptotic cells at the early timepoint, whereas as the majority of aggregates become hyperconfluent and self-detached an increasing proportion exhibit apoptosis (n = 53 aggregates).

As cytosolic YAP is not observed in sparse regions with low prevalence of apoptosis, we sought to determine if YAP activation (i.e., translocation to the nucleus) is sufficient to inhibit apoptosis. We stably expressed activated YAP in VICs and cultured them on circular collagen islands until hyperconfluent (but prior to self-detachment). Compared to control VICs (Fig. 10A), we observe a dramatic decrease in apoptotic activity in VICs with constitutively active YAP (Fig. 10B).

**Figure. 10.**
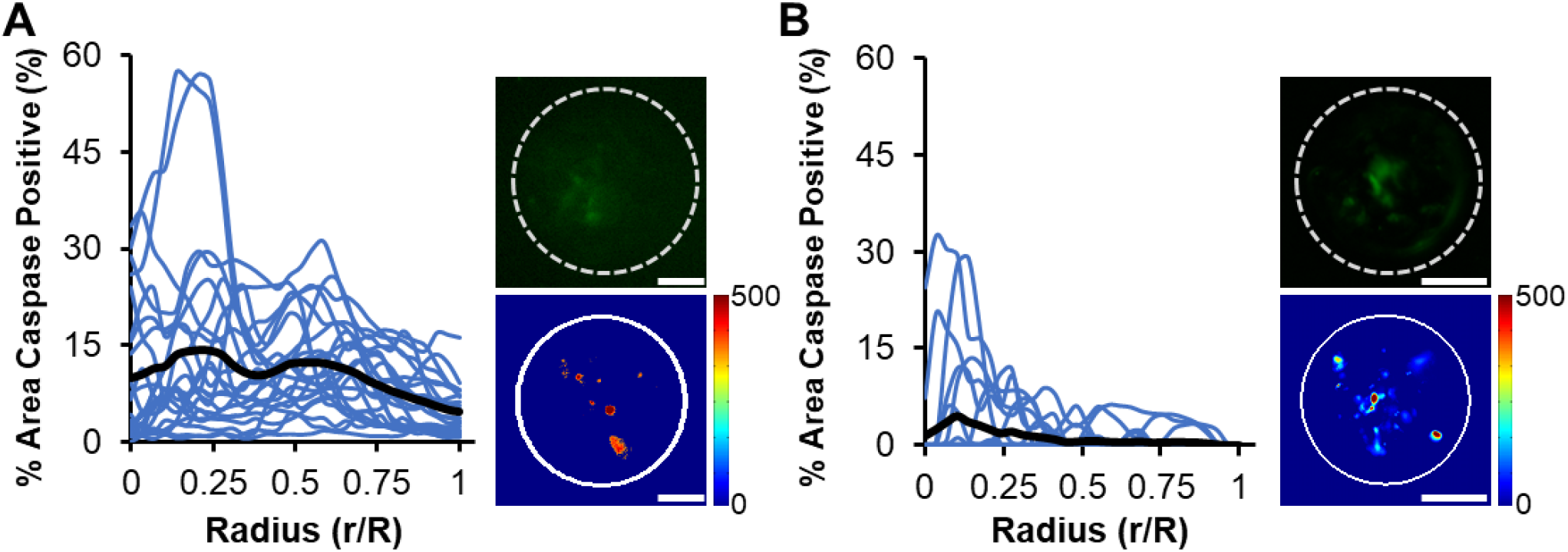
Apoptosis is decreased substantially by YAP activation. A) The portion of aggregates positive for caspase in control VICs decreases from the center to edge. Image of a single aggregate stained for cleaved caspase-3/7 (top inset; green, dotted line indicates aggregate edge) and heat map for average caspase-3/7 intensity (bottom inset, n = 24 aggregates from 3 replicates). B) The portion of aggregates positive for caspase in VICs with constitutively active YAP decreases from center to edge. Image of a single aggregate stained for cleaved caspase-3/7 (top inset; green, dotted line indicates aggregate edge) and heat map for average caspase-3/7 intensity (bottom inset, n = 23 aggregates from 3 replicates). Scale bar = 50 μm.

## 4. Discussion

Collective cell interactions drive distinct spatial patterning of cell behavior in monolayers of cells within confined geometries. These emergent patterns correlate strongly with predicted mechanical stress fields indicating a regulatory role of mechanics. Here, we extended studies conducted primarily on epithelial monolayers to more contractile and less contact inhibited primary fibroblastic cells. We analyzed the evolution of VIC confluency on circular micropatterned protein islands over a period of days, a long duration which is relevant for disease modeling. We aimed to determine how observed heterogeneities in spatial patterning are influenced by interactions between neighboring cells and under the influence of the circular global constraint to determine the relative effects of local and global collective behaviors. In contrast to previous reports of radially symmetric cell alignment, proliferation, and differentiation of epithelial cells and cell lines in circular and annular patterns (Nelson et al. 2005, Streichan et al. 2014, He et al. 2015, Silver et al. 2020), we observe that primary porcine VICs and human dermal fibroblasts align locally and form hyperconfluent bands spanning the circular patterns due to local cell-cell interactions, and this local alignment occurs regardless of pattern diameter (200-600 μm). The banding behavior bears similarity to local collective behaviors reported in unconstrained fibroblast monolayers from a variety of sources (Carlson et al. 2009, Lynch et al. 2018, Chansard et al. 2021). The asymmetric locations of bands largely explain the variability in patterns of proliferation, migration, and density observed in these aggregates. The formation of these highly aligned multicellular bands soon after seeding accelerates the rate to confluency resulting in aggregate-to-aggregate heterogeneity within a single dish of identical collagen islands. These observations, along with traction force and time-lapse migration measurements, indicate that VICs exhibit strong local collective behavior that disrupts the emergent behavior emanating from the circular constraint of the islands. Such regional changes in cell shape, number, and confluency should be considered in the mechanical analysis of multicellular systems, especially for primary fibroblastic cells.

Non-uniform stress fields that emerge from transmission of forces between cells have been postulated as the driver of cell alignment and collective behavior in constrained cell islands (Nelson et al. 2005, Li et al. 2009, Silver et al. 2020). In contrast, our short-term (4.5 hr) time-lapse imaging shows that the cell alignment that drives the formation of bands is initiated by a few local cells migrating to and aligning with a single highly elongated spindle-shaped cell. Rather than acting individually until confluent as seen in epithelial cells (Doxzen et al. 2013), VICs often align with each other within the patterns at the pre-confluent stage. This local collective behavior may occur due to cells sensing the anisotropic displacement of the underlying PA gel generated by the cell (Reinhart-King et al. 2005) and/or by contact guidance provided by the elongated cell. This behavior occurs regardless of aggregate size up to 600 μm diameter indicating that the banding is not formed due edge effects, i.e., cells do not attach to two points on the circular edge and elongate to span the island. We show that the banding also occurs with primary dermal fibroblasts. Sun and colleagues report that primary rat embryonic fibroblasts align radially at the edge of similar protein islands, and the central cells are smaller and more dense than edge cells, although without banding (Xie et al. 2021). This heterogeneous morphology is stark contrast to the patterns observed for NIH3T3 and osteoblast-like MC3T3-E1 cell lines which exhibit uniform spread area throughout the constrained monolayers and circumferentially aligned cells at the edges and less elongated cells in the center (He et al. 2015, Xie et al. 2021). Together, these results indicate that for the relatively large and contractile primary fibroblastic cells studied herein, the strong local collective behavior disrupts the global circular constraint that drives the radially symmetric emergent collective behavior of other cell types cultured on circular patterns.

The long-term (24 hr) time-lapse videos clearly show that even when hyperconfluent, confined VIC cell layers are not static or “jammed” as observed for high-density epithelial monolayers (Garcia et al. 2015, Vig et al. 2017, Atia et al. 2018). We observe synchronized circumferential collective migration at the edges of pre-confluent and confluent VIC aggregates as well as substantial collective movement in the center along the multicellular bands. Similar collective cell migration is observed in epithelial monolayers on small (100-200 μm) circular islands, and vortices of local collective migration are observed in the central regions of larger islands (500-1000 μm) (Doxzen et al. 2013). The average cell speed was not significantly lower in hyperconfluent regions than in less confluent regions. Although this result is not consistent with the common finding that collective cell migration speed has an inverse correlation with cell density (Doxzen et al. 2013, Li et al. 2014, Lin et al. 2021), the motion of the cells in the bands appears qualitatively different in that it is less persistent in direction and more fluctuating.

The progression of bands to hyperconfluency appears to be driven by cells migration to and joining aggregates rather than proliferation within the bands. Both EdU staining and cell morphological features in time-lapse videos show that proliferation occurs throughout the aggregates except within the hyperconfluent regions. Previous experiments with long EdU pulses (8 hr) show EdU staining throughout similar constrained monolayers (mouse MSC cell line) with a decrease in proliferation with cell density (Berent et al. 2022). In contrast, many studies of constrained epithelial monolayers report proliferation occurring predominantly at the edges corresponding to strong cell alignment and high traction forces (Li et al. 2006, Streichan et al. 2014, Silver et al. 2020). Utilization of low Ca^2+^ media, shown to disrupt cell-cell interactions and subsequently the traction and cell alignment at the edges of cell aggregates (Maruthamuthu et al. 2011, Xie et al. 2021), did not inhibit the alignment of cells into bands across our aggregates but did stop the bands from progressing to hyperconfluency. These results indicate that development of the hyperconfluent bands is dependent upon cell-cell force transmission; however, the results should be interpreted carefully as the initial cell alignment occurred in the 48 hours prior to depleting calcium from media (Fig. 4A), and decreased proliferation rates in low-calcium media may contribute to the lower cell density (Kahl et al. 2003). Regardless, the results demonstrate that removing calcium from the media does not reverse the local collective behavior or fully block VIC proliferation.

The hyperconfluent bands produce high, localized traction stresses at their endpoints. In contrast, in previous studies with confluent islands of epithelial cells and cell lines that are not highly motile (Schaumann et al. 2018), traction stresses are highest and roughly uniform at the edges, and the predicted cell-layer stresses increase from the edge to the center (Nelson et al. 2005, Li et al. 2009, Aragona et al. 2013, Deglincerti et al. 2016, Tran et al. 2020). Time-lapse TFM heatmaps demonstrate that the magnitudes of traction stress vary with time but are consistently located at band endpoints and are directed inward indicating high uniaxial tension in bands. Traction stresses increase as bands progress from confluent to hyperconfluent and eventually lead to the detachment of portions of the aggregates from the substrate when the collagen/substrate bond strength is exceeded. Paradoxically, despite high traction forces at band ends, low-stress markers are observed in cells in the bands. Cells in hyperconfluent regions have smaller areas and cytosolic YAP; markers that are characteristic of cells in low-stress environments such as when cultured on soft substrates or on small protein islands that constrain spreading (Wada et al. 2011, Aragona et al. 2013, Calvo et al. 2013, Mascharak et al. 2017). In contrast, cells at band endpoints have more nucleated YAP similar to that observed for single cells on unpatterned stiff substrates (Dupont et al. 2011) and for confluent primary fibroblasts on protein islands (Xie et al. 2021). Apoptosis, which is also associated with low cell stress (Egerbacher et al. 2008, Humphrey et al. 2014), increases as aggregates progress to hyperconfluency. Consistent with this result, almost every aggregate that can no longer generate substantial traction due to being partially detached from the substrate is caspase positive. As YAP enhances the transcription of pro-survival genes (Zhang et al. 2011, Codelia et al. 2014, Lin et al. 2015), we wondered if YAP plays a defining role in apoptosis in VICs. We found that constitutively activating YAP substantially decreases apoptosis consistent with findings from studies of epithelial cell lines (Liu et al. 2017). While high traction stresses measured at the ends of the bands seem contradictory with low stress within the bands, it is possible that the high force in the band is generated/transmitted by many cells in parallel, i.e., high total force but low force per cell. This interpretation is consistent with our computational model of circular cell aggregates with radially increasing contractility from the center to the edge which predicts low stress in the central region despite high traction stresses transmitted to the substrate at the edges (Goldblatt et al. 2020).

To further investigate the state of stress in the bands, we applied cyclic uniaxial stretch to confluent aggregates. Under the assumption that cells attempt to reach a homeostatic stress level (Brown et al. 1998), we postulated that if cells within the bands were under high tension, they would reorient away from stretch to regain their preferred stress level, whereas if cells were confined due to high cell density and not able to generate their preferred level of homeostatic tension, they would spread out and reorient towards stretch. Isolated VICs (Cirka et al. 2016) and other fibroblasts cultured on stiff substrates exhibit strain avoidance (Wang et al. 2001, Greiner et al. 2013, Shao et al. 2013, Ristori et al. 2018) as do epithelial/endothelial sheets (Kaunas et al. 2005, Gérémie et al. 2022). In contrast, when cells are cultured on collagen gels (Tondon et al. 2014), within low density soft collagen gels (Foolen et al. 2014, Sears et al. 2016) or when contractility is inhibited (Kaunas et al. 2005), cells reorient towards stretch. Consistent with the low stress interpretation, we observed that VICs within bands that are already oriented towards the direction of stretch elongate and have more pronounced and aligned cytoskeletal arrangement, whereas cells within bands roughly perpendicular to stretch retracted or reoriented towards the direction of stretch. We did not observe increased aggregate detachment with stretch which also suggests low stress in the aggregates. Cyclic (equibiaxial) stretch of VIC monolayers, in combination with TGF-beta treatment, has been shown to potentiate cell detachment and formation of aggregates (Fisher et al. 2012). As we presumed that edge cells were under relatively high circumferential stress due to their elongated morphology and based on computational model predictions of high circumferential stresses at the edges of constrained monolayers (He et al. 2015), we expected these cells to reorient away from the stretch direction. Instead, we observed that the cells which were parallel to the direction of stretch maintained their elongation, and cells that were perpendicular to the direction of stretch retracted and/or reoriented away from the stretch direction. Cyclically stretched epithelial (MDCK) cells have been shown to increase their polarization (aspect ratio) when aligned around a “wound” edge in a monolayer model with a hole (essentially the inverse of our “island” system) consistent with the behavior we observe for cells aligned with the stretch axis, but the retraction of edge cells perpendicular to stretch has not been reported (Xu et al. 2022). It is possible that the aligned bands in our system disrupt the emergent radial-symmetric stress field resulting in relatively low stress in edge cells, thus these cells reorient in the stretch direction to increase their stress towards a homeostatic level or to minimize shear stress (Liu et al. 2016).

## 5. Conclusion

Our results indicate that cell alignment and subsequent local collective behavior leads to substantial heterogeneity between and within constrained monolayers of primary fibroblastic cells. Multicellular bands are formed by cell-cell interactions locally. As cells become more dense within the bands, the stress exerted by interior cells decrease, YAP becomes excluded from the nucleus and, consequently, apoptotic rates increase. These findings suggest that strong local cell-cell interactions between primary fibroblastic cells can disrupt the global collective cellular behavior; this local collective cell behavior may play an important role in development and disease of connective tissues.

## Acknowledgment

We would like to thank Sabine Hahn for her help editing the manuscript, Seyed Mohammad Jafar Sobhani for his help with graphical plotting with MATLAB, as well as Marsha Rolle and CERES labs located in Worcester Polytechnic Institute for use of the Cytation microscope and George Pins for donation of neonatal human dermal fibroblastic cells. This work was funded in part by the American Heart Association (20AIREA35120448) and the National Science Foundation (CMMI 1761432).

