## Supplemental Figures for "Multicellular Aligned Bands Disrupt Global Collective Cell Behavior"


### Graphical Abstract

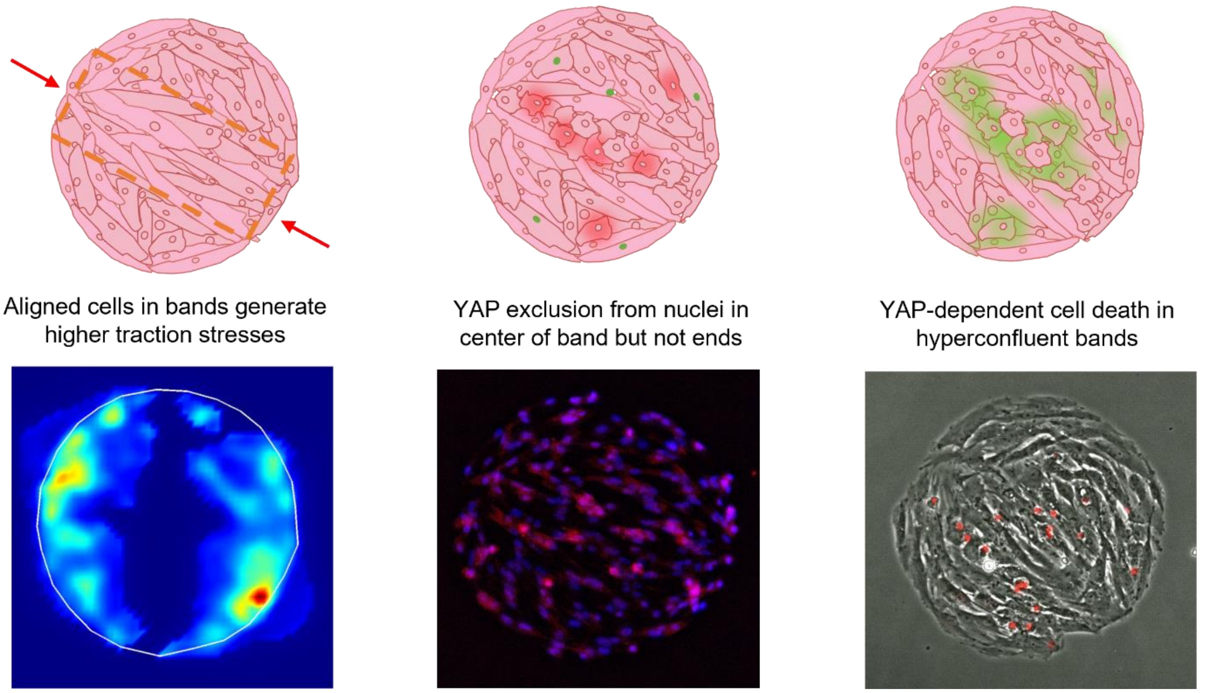

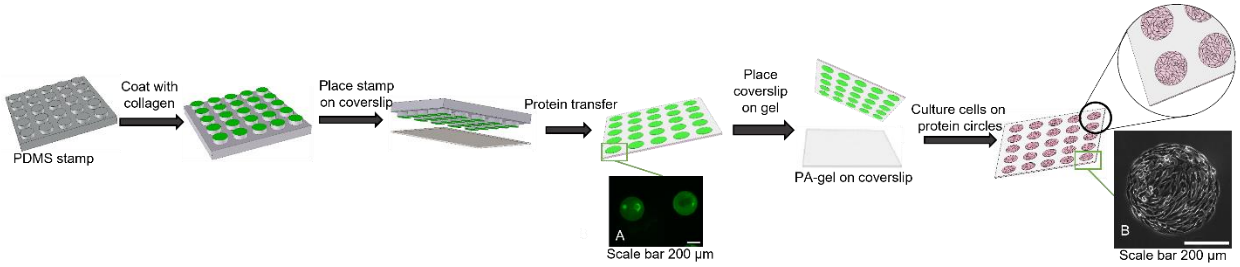


*Figure S.1. Indirect microcontact printing method schematic, modified from [26]. Inset (A) Collagen prints verified by Alexa Fluor-488 carboxylic acid succinimidyl ester. The uniform distribution of the collagen is shown in the indirect microcontact printing method used in this research. Inset (B) Pre-confluent 2D aggregate on 400 µm in diameter collagen island showing cell coverage and circular shape of the print. The scale bar is 200 µm.*

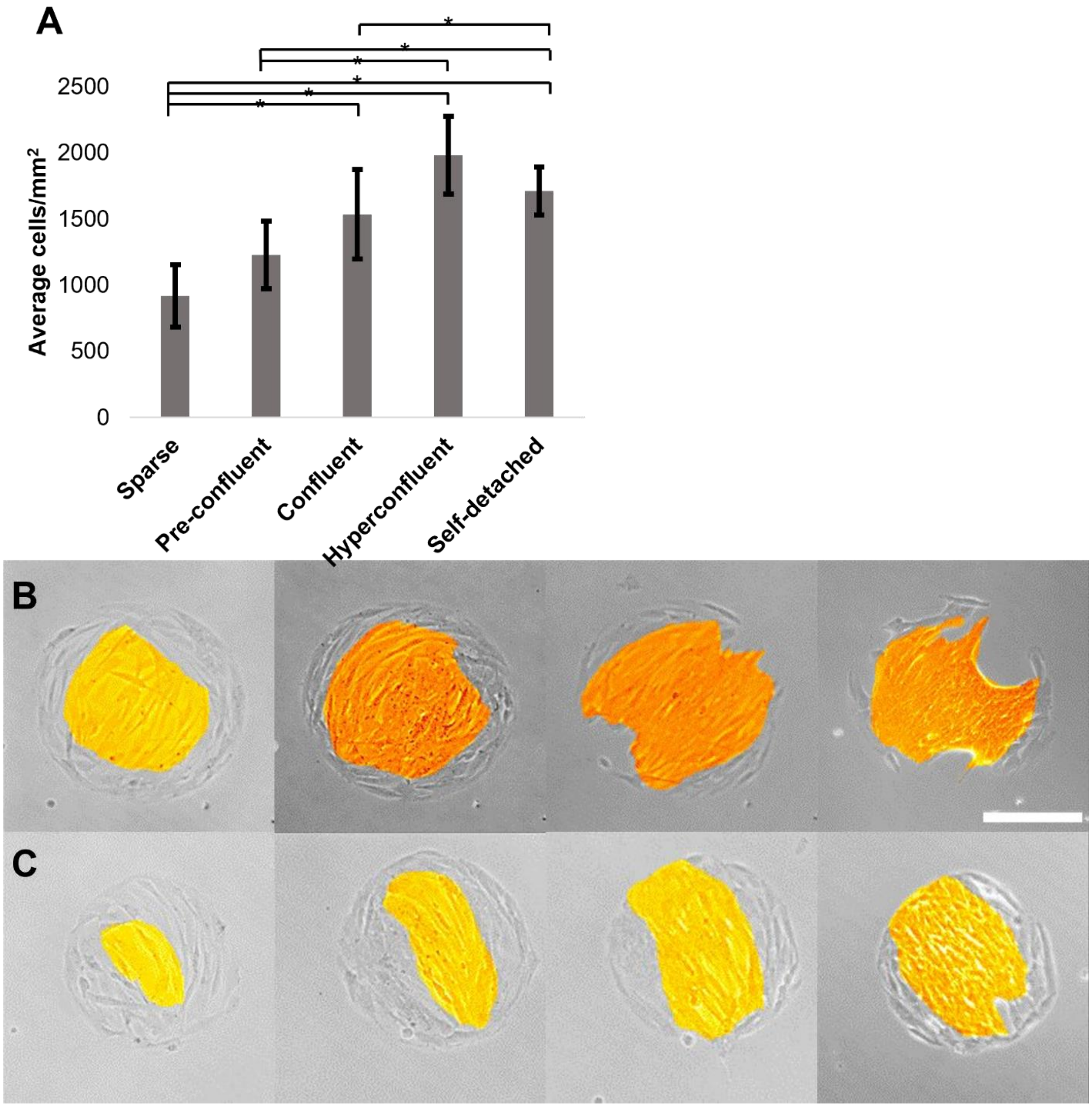


*Figure S.2. The cell density increases with confluency level (A), but the confluency progress differently in each aggregate (B, C). Adjacent confluency levels are qualitatively different but do not have significantly different cell density due to heterogeneity in each aggregate (ANOVA, * indicates p<0.05). Aggregate in (B) developed alignment in pre-confluent level and progressed quickly to self-detachment. Aggregate in (C) has local alignment only in small region in pre-confluent level and progressed slowly hype confluency and shows only minor self-detachment. Yellow/orange highlight indicates aligned region. Scale bar = 200 µm.*

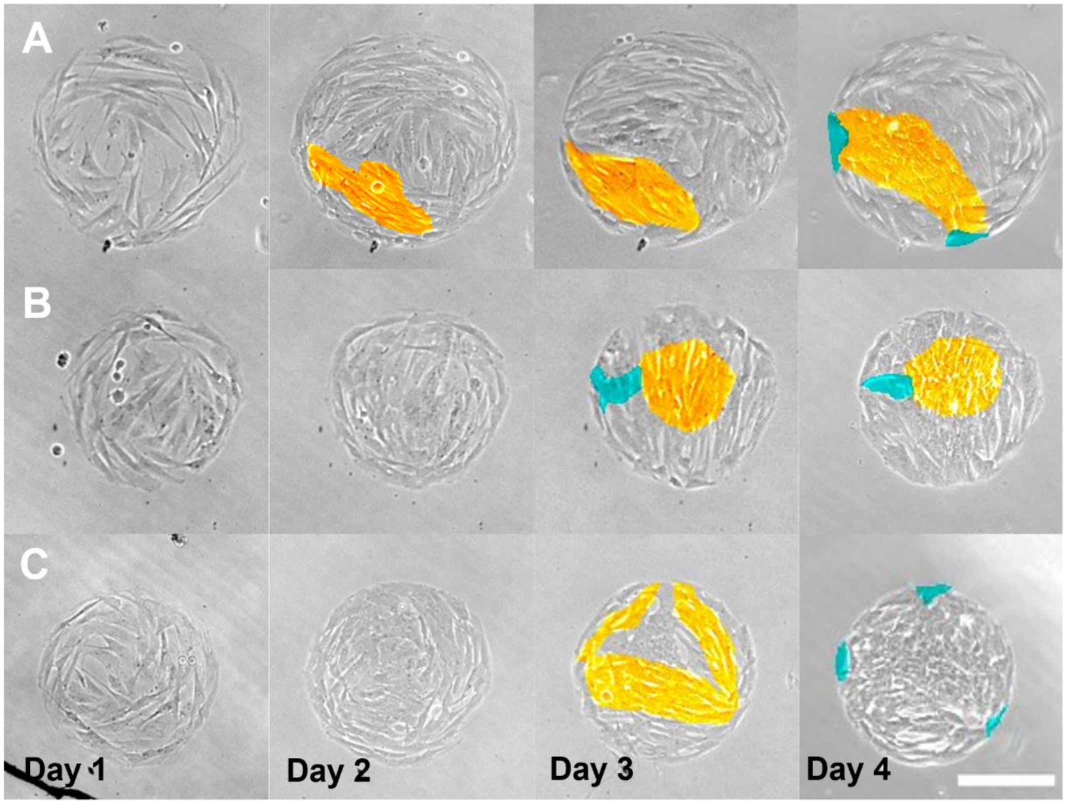


*Figure S.3. Local collective behavior results in hyperconfluent bands of many shapes. A) Aggregate with banding initiating non-centrally. B) Aggregate with central banding that never spans the entire island to meet the edge, and hole appears in low density region. C) Aggregate with banding in three directions forming a triangular pattern and detachment in three locations. Yellow/orange highlight indicates aligned region; blue highlight indicates detachment of the cells from the substrate. Scale bar = 200 µm.*

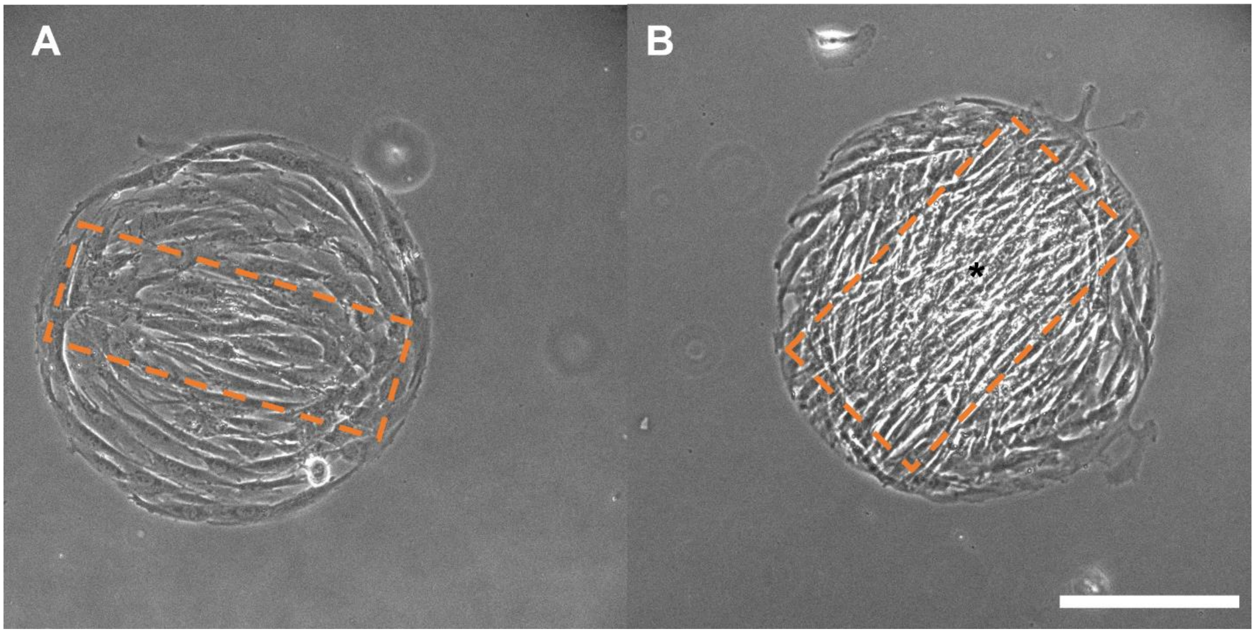


*Figure S.4. Banding (A, B) and hyperconfluency (B) are observed in human dermal fibroblast (HDF) aggregates (* indicates hyperconfluent region). Scale bar = 200 µm.*

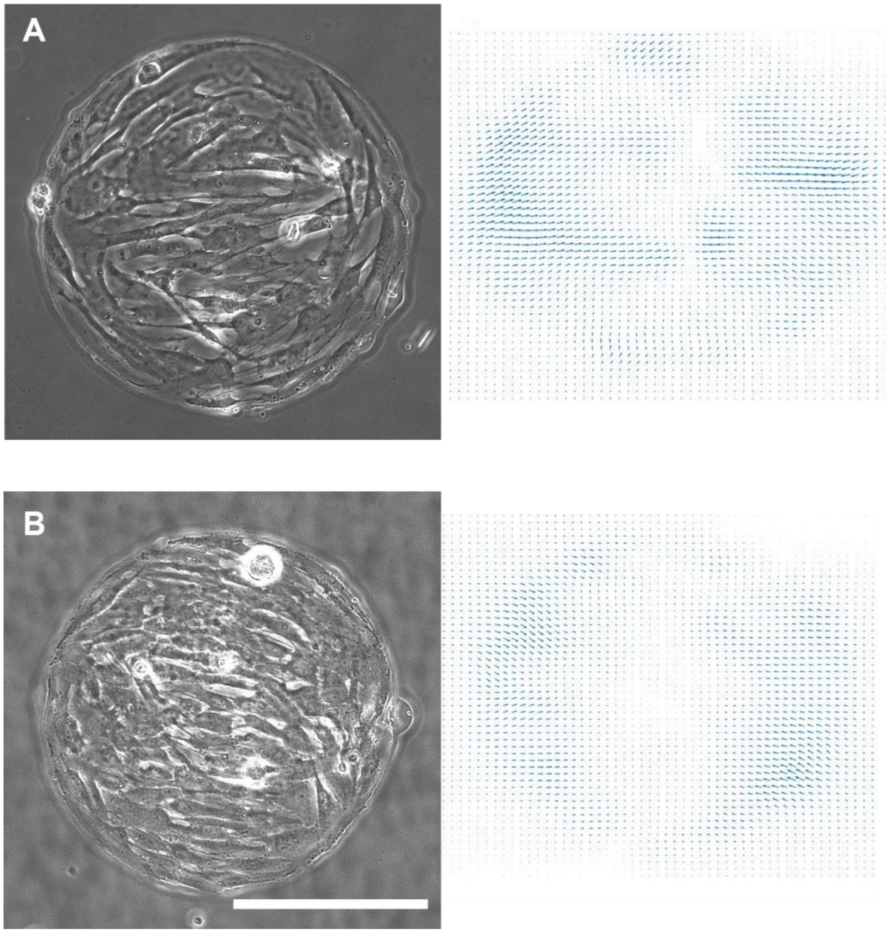


*Figure S.5. TFM vector plots show direction of traction force is along the long axis of the band. A) Phase image of a sparse aggregate and corresponding traction force vectors. B) Phase image of a confluent aggregate and corresponding traction force vectors. Scale bar = 200 μm.*

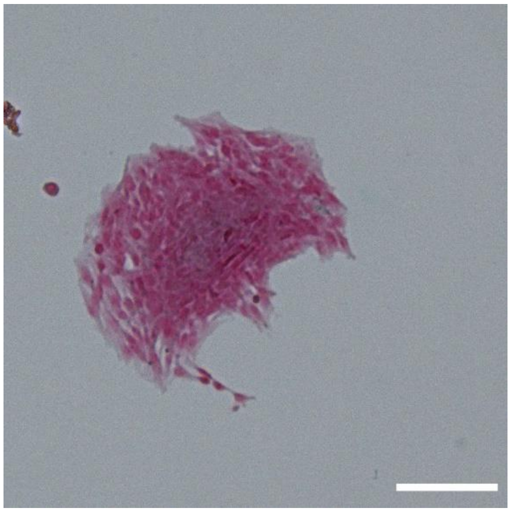

*Figure S.6. Collagen stained after self-detached stage shows the removal of the collagen from the PA gel surface. Scale bar = 200 μm. The brightness of images has been manipulated to purpose of presentation.*

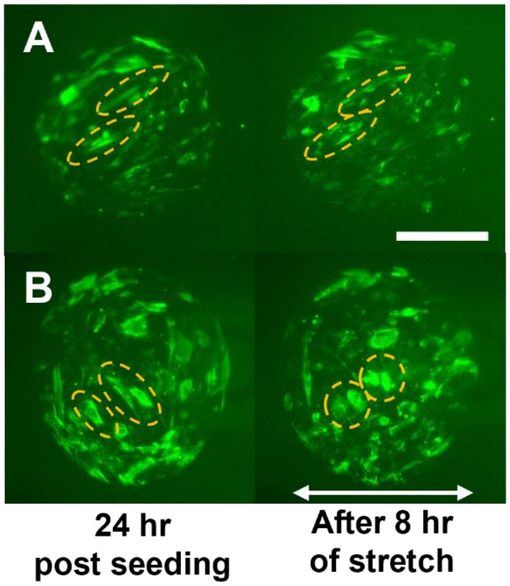


*Figure S.5. Dynamic stretch reinforces F-actin alignment for cells in the direction of stretch and disrupts the cytoskeleton of cells aligned away from the stretch axis. A) Representative example of cells within a band aligned at an intermediate angle (~30°) in which the F-actin is intensified but the cell reorientation is minimal. B) Representative example of elongated cells within a band oriented greater than 45° from the stretch axis in which the F-actin cytoskeleton is disrupted and the cells become rounded. For all panels, F-actin of live VICs was stained with CellMask™ Actin before and after application of 10% uniaxial stretch for 8 hr at 1 Hz; brightly stained cells inside of bands are outlined by ellipses to show change in orientation and elongation. Scale bar = 200 µm.*
