## Supplemental Movie 2 for "Multicellular Aligned Bands Disrupt Global Collective Cell Behavior"

### Slide 1
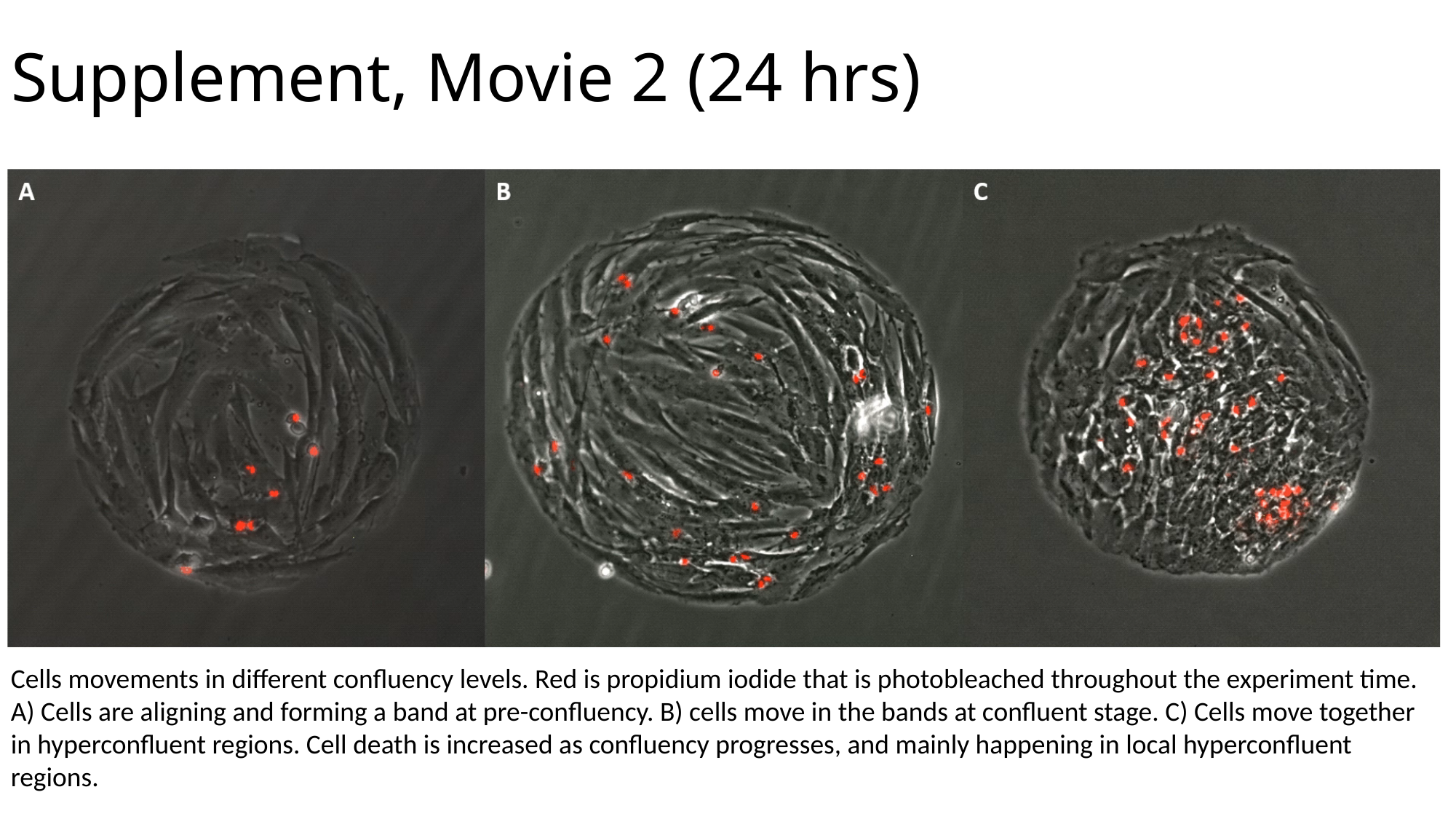

Supplement, Movie 2 (24 hrs)
Cells movements in different confluency levels. Red is propidium iodide that is photobleached throughout the experiment time. A) Cells are aligning and forming a band at pre-confluency. B) cells move in the bands at confluent stage. C) Cells move together in hyperconfluent regions. Cell death is increased as confluency progresses, and mainly happening in local hyperconfluent regions.
