## Supplemental Movie 1 for "Multicellular Aligned Bands Disrupt Global Collective Cell Behavior"

### Slide 1
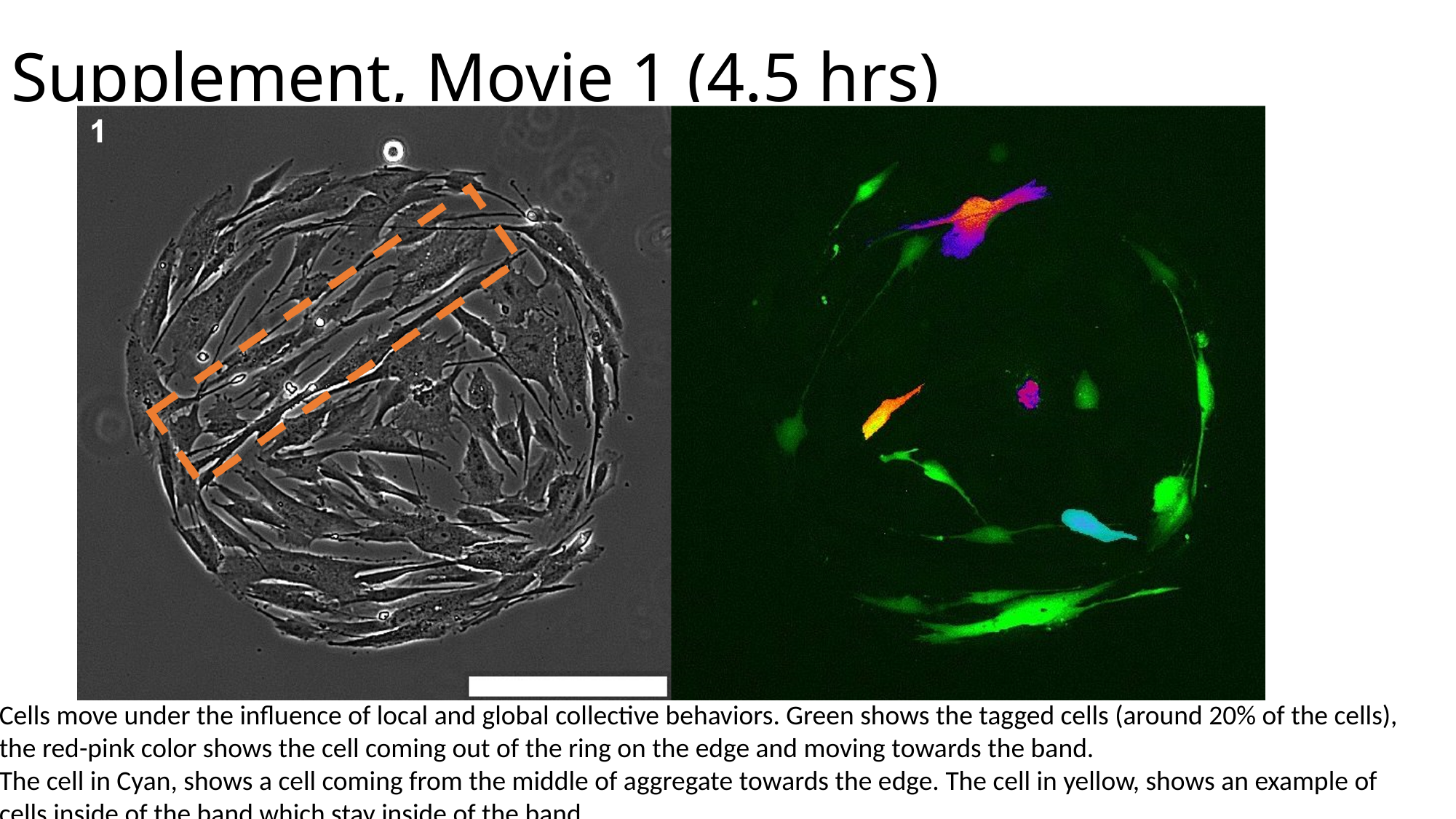

Supplement, Movie 1 (4.5 hrs)
Cells move under the influence of local and global collective behaviors. Green shows the tagged cells (around 20% of the cells), the red-pink color shows the cell coming out of the ring on the edge and moving towards the band.
The cell in Cyan, shows a cell coming from the middle of aggregate towards the edge. The cell in yellow, shows an example of cells inside of the band which stay inside of the band.
