## Supplemental Movie 3 for "Multicellular Aligned Bands Disrupt Global Collective Cell Behavior"

### Slide 1
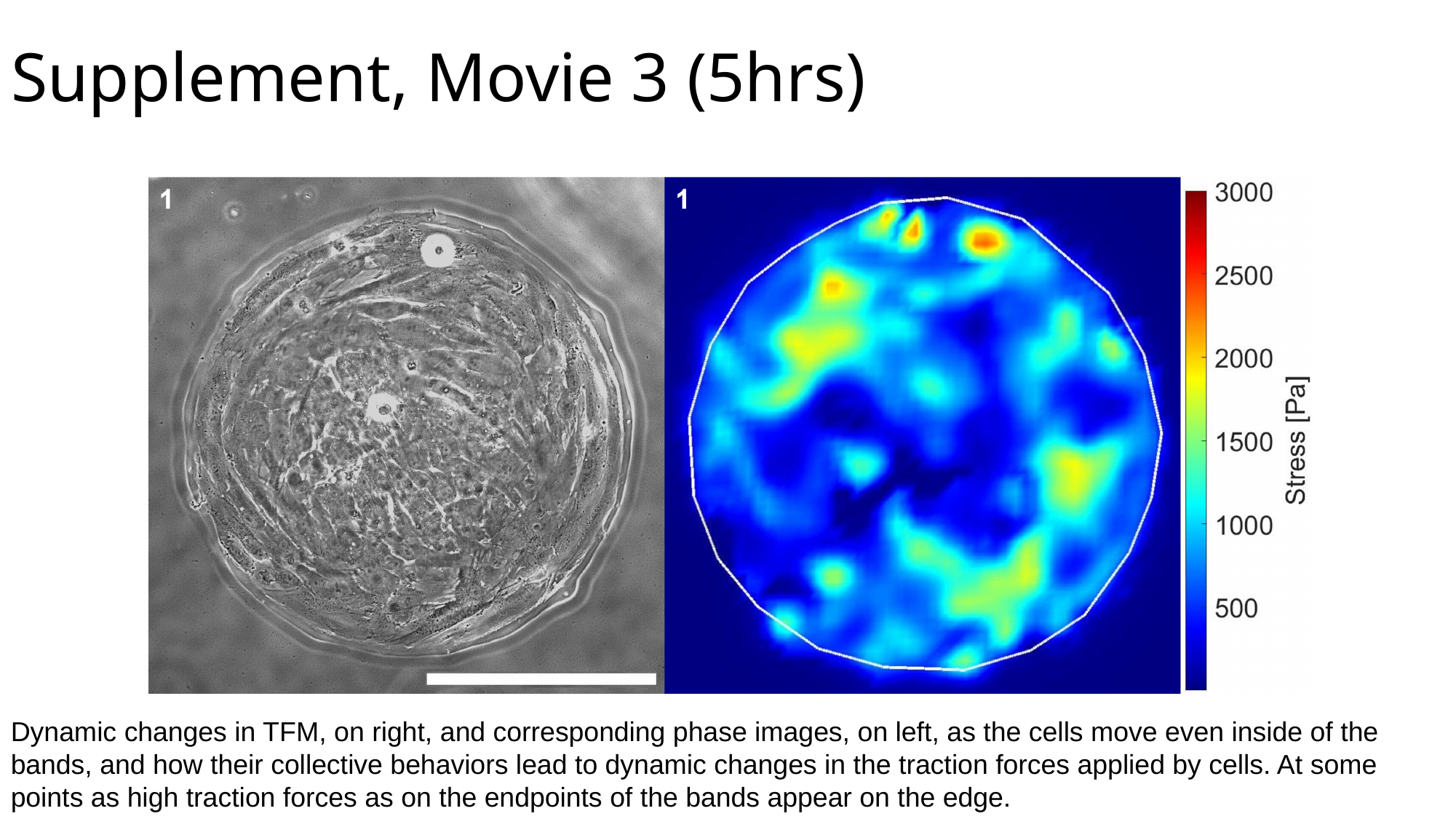

Supplement, Movie 3 (5hrs)
Dynamic changes in TFM, on right, and corresponding phase images, on left, as the cells move even inside of the bands, and how their collective behaviors lead to dynamic changes in the traction forces applied by cells. At some points as high traction forces as on the endpoints of the bands appear on the edge.
